## Supplemental Dataset 3 for "Ptbp1 is not required for retinal neurogenesis and cell fate specification"

### Extended Dataset 3

#### A. Rod Specific Exons

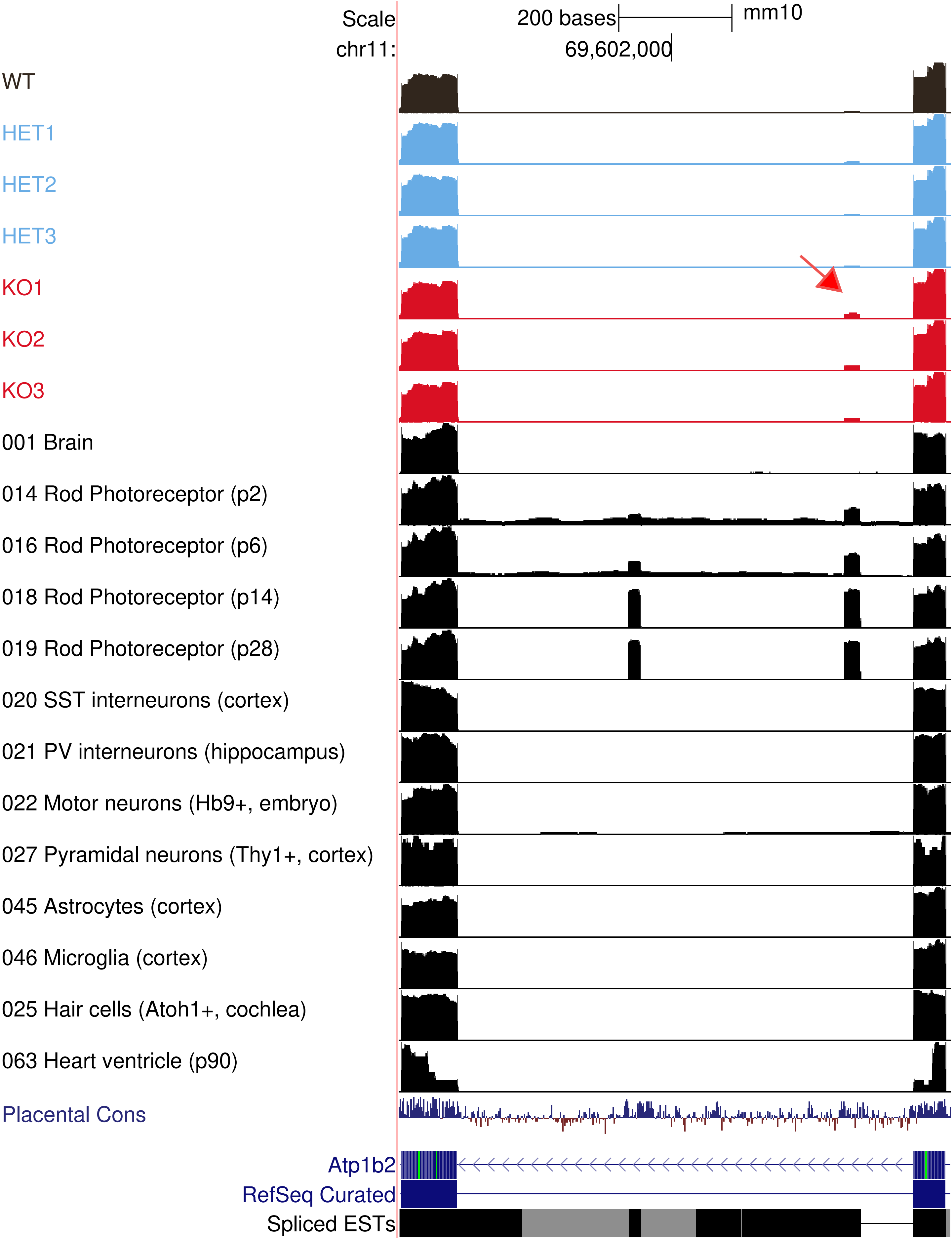

Scale
200 bases
mm10
chr11:
69,602,000
WT
HET1
HET2
HET3
KO1
KO2
KO3
001 Brain
014 Rod Photoreceptor (p2)
016 Rod Photoreceptor (p6)
018 Rod Photoreceptor (p14)
019 Rod Photoreceptor (p28)
020 SST interneurons (cortex)
021 PV interneurons (hippocampus)
022 Motor neurons (Hb9+, embryo)
027 Pyramidal neurons (Thy1+, cortex)
045 Astrocytes (cortex)
046 Microglia (cortex)
025 Hair cells (Atoh1+, cochlea)
063 Heart ventricle (p90)
Placental Cons
Atp1b2
RefSeq Curated
Spliced ESTs

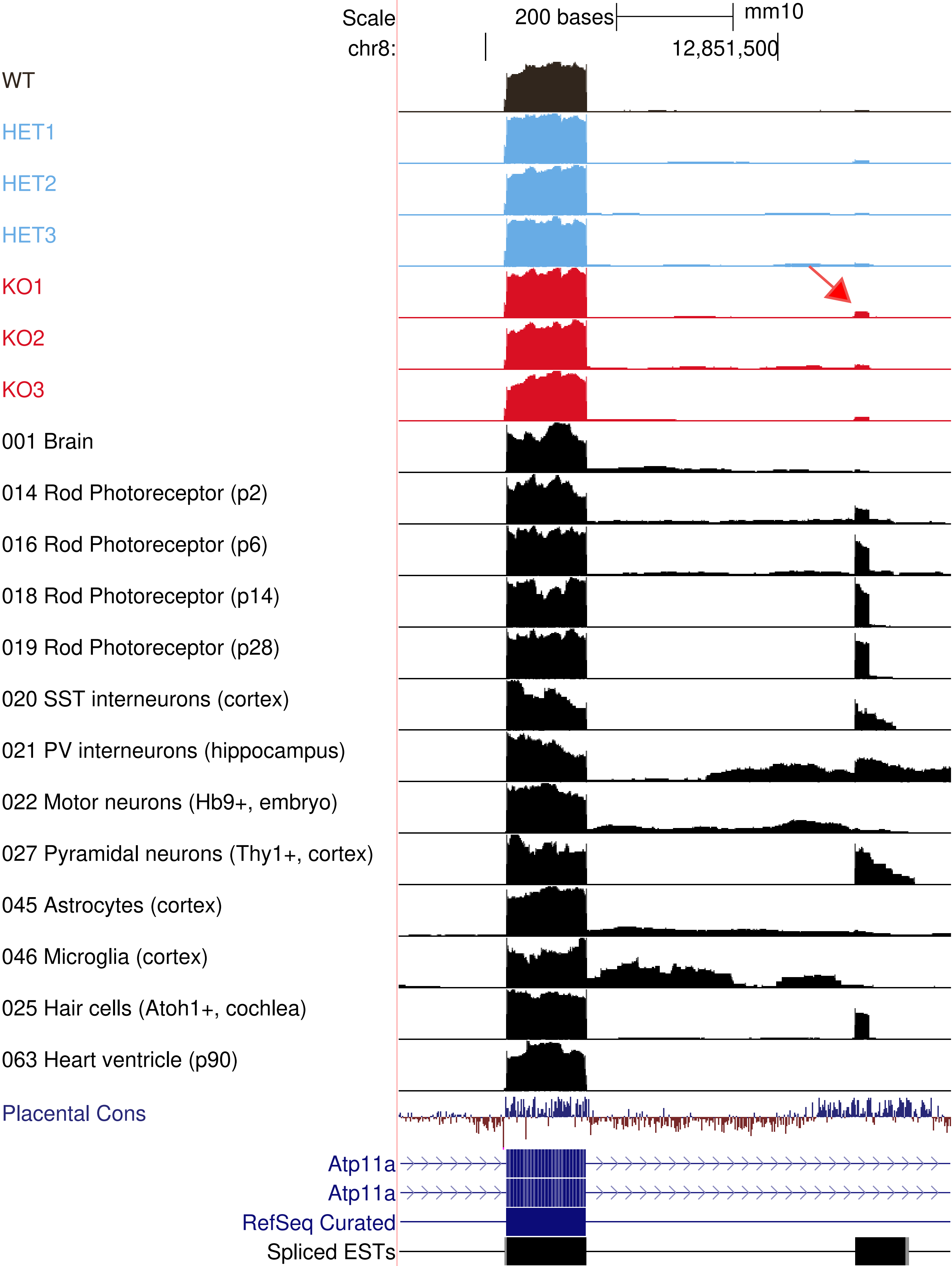

Scale
200 bases
mm10
chr8:
12,851,500
WT
HET1
HET2
HET3
KO1
KO2
KO3
001 Brain
014 Rod Photoreceptor (p2)
016 Rod Photoreceptor (p6)
018 Rod Photoreceptor (p14)
019 Rod Photoreceptor (p28)
020 SST interneurons (cortex)
021 PV interneurons (hippocampus)
022 Motor neurons (Hb9+, embryo)
027 Pyramidal neurons (Thy1+, cortex)
045 Astrocytes (cortex)
046 Microglia (cortex)
025 Hair cells (Atoh1+, cochlea)
063 Heart ventricle (p90)
Placental Cons
Atp11a
Atp11a
RefSeq Curated
Spliced ESTs

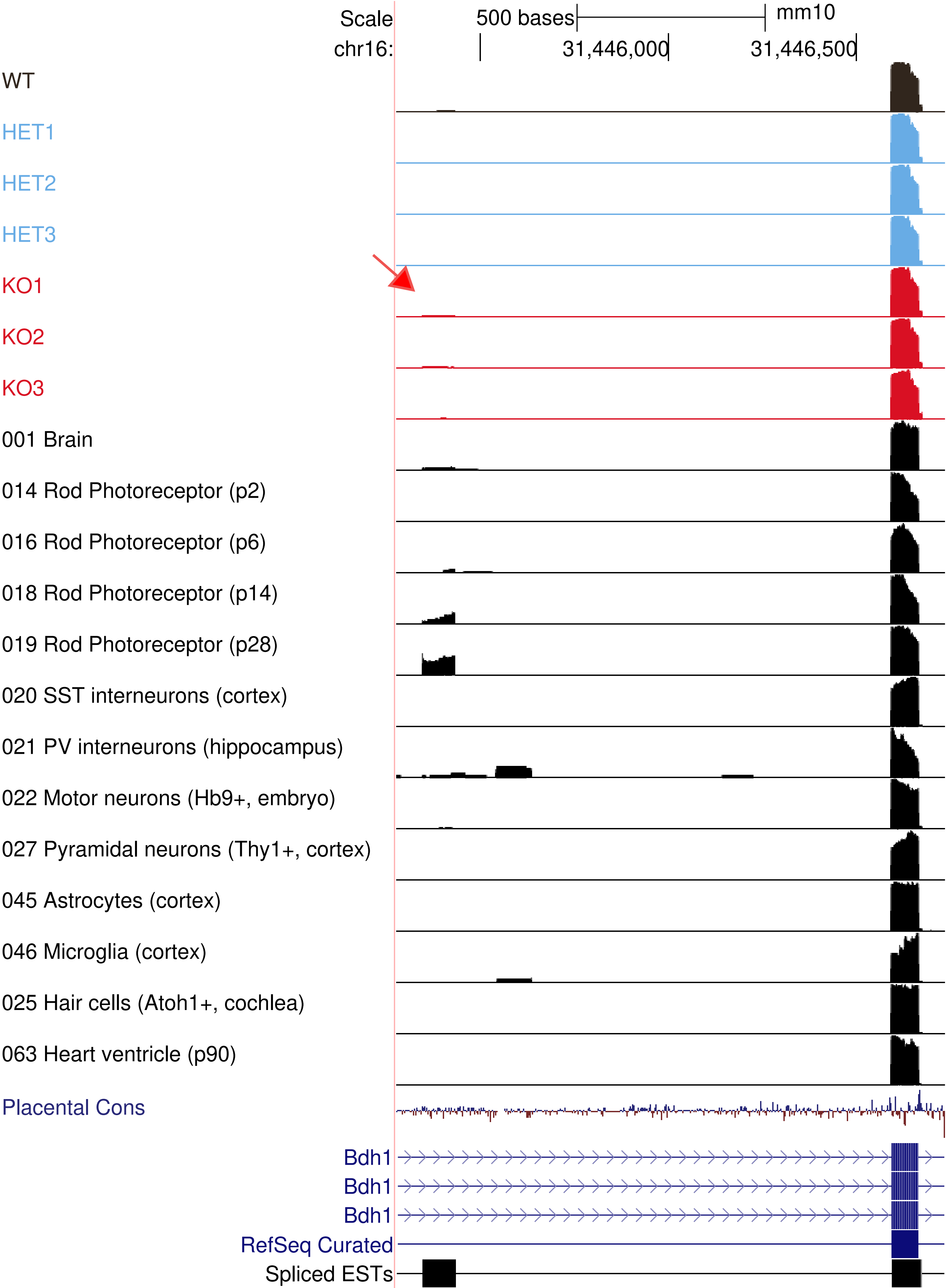

Scale
500 bases
mm10
chr16:
31,446,000
31,446,500
WT
HET1
HET2
HET3
KO1
KO2
KO3
001 Brain
014 Rod Photoreceptor (p2)
016 Rod Photoreceptor (p6)
018 Rod Photoreceptor (p14)
019 Rod Photoreceptor (p28)
020 SST interneurons (cortex)
021 PV interneurons (hippocampus)
022 Motor neurons (Hb9+, embryo)
027 Pyramidal neurons (Thy1+, cortex)
045 Astrocytes (cortex)
046 Microglia (cortex)
025 Hair cells (Atoh1+, cochlea)
063 Heart ventricle (p90)
Placental Cons
Bdh1
Bdh1
Bdh1
RefSeq Curated
Spliced ESTs

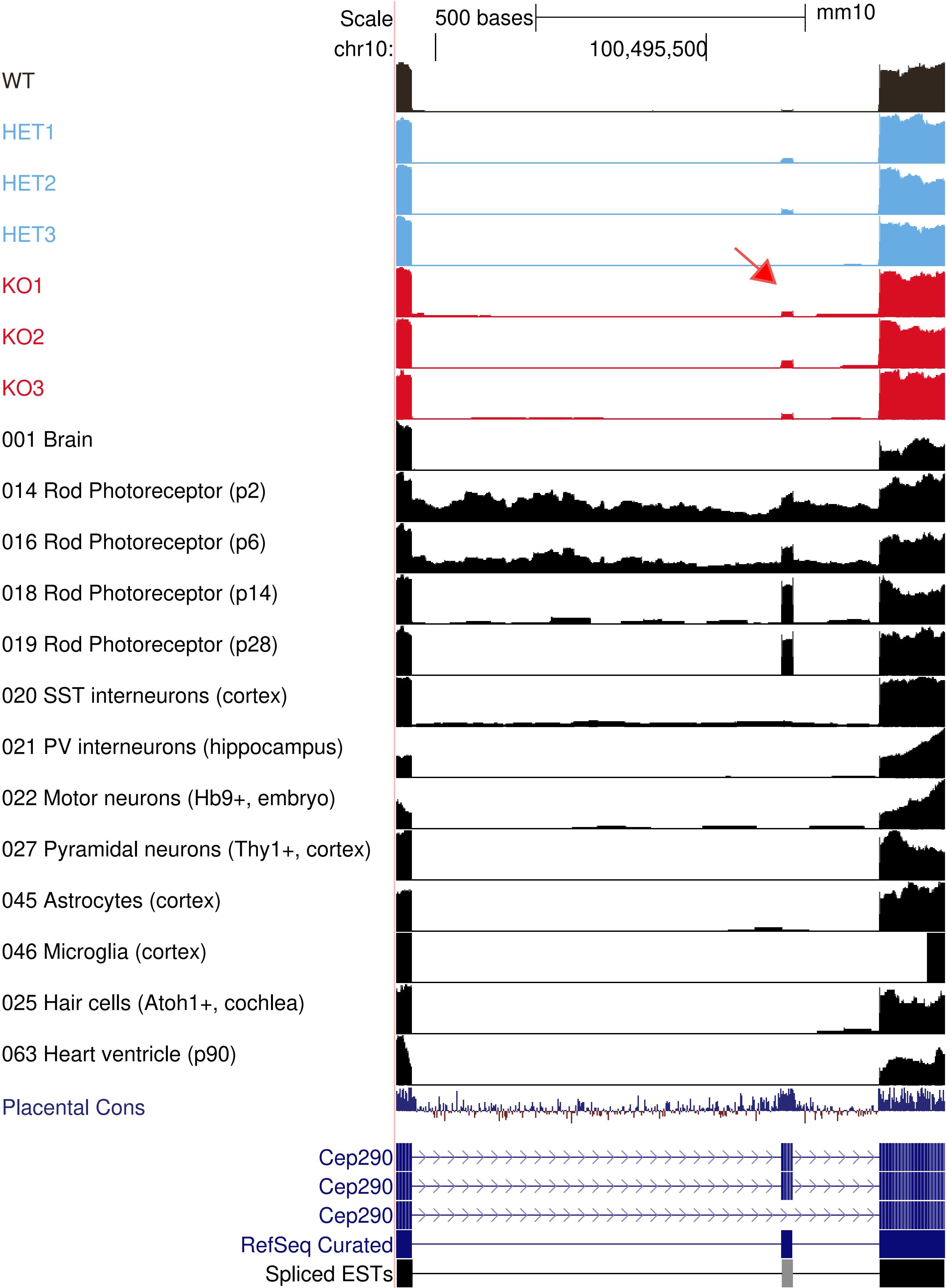

Scale
500 bases
mm10
chr10:
100,495,500
WT
HET1
HET2
HET3
KO1
KO2
KO3
001 Brain
014 Rod Photoreceptor (p2)
016 Rod Photoreceptor (p6)
018 Rod Photoreceptor (p14)
019 Rod Photoreceptor (p28)
020 SST interneurons (cortex)
021 PV interneurons (hippocampus)
022 Motor neurons (Hb9+, embryo)
027 Pyramidal neurons (Thy1+, cortex)
045 Astrocytes (cortex)
046 Microglia (cortex)
025 Hair cells (Atoh1+, cochlea)
063 Heart ventricle (p90)
Placental Cons
Cep290
Cep290
Cep290
RefSeq Curated
Spliced ESTs

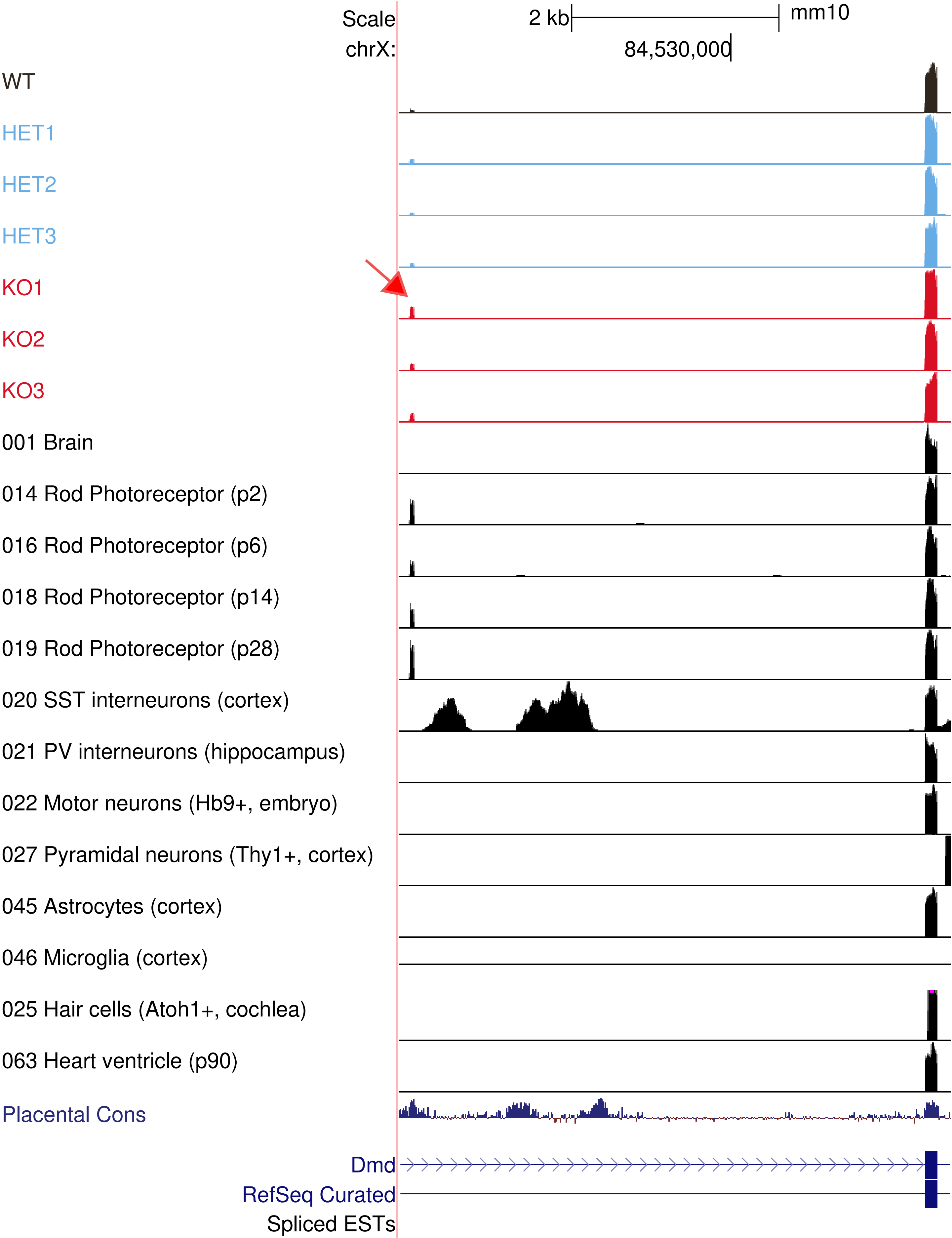

Scale
2 kb
mm10
chrX:
84,530,000
WT
HET1
HET2
HET3
KO1
KO2
KO3
001 Brain
014 Rod Photoreceptor (p2)
016 Rod Photoreceptor (p6)
018 Rod Photoreceptor (p14)
019 Rod Photoreceptor (p28)
020 SST interneurons (cortex)
021 PV interneurons (hippocampus)
022 Motor neurons (Hb9+, embryo)
027 Pyramidal neurons (Thy1+, cortex)
045 Astrocytes (cortex)
046 Microglia (cortex)
025 Hair cells (Atoh1+, cochlea)
063 Heart ventricle (p90)
Placental Cons
Dmd
RefSeq Curated
Spliced ESTs

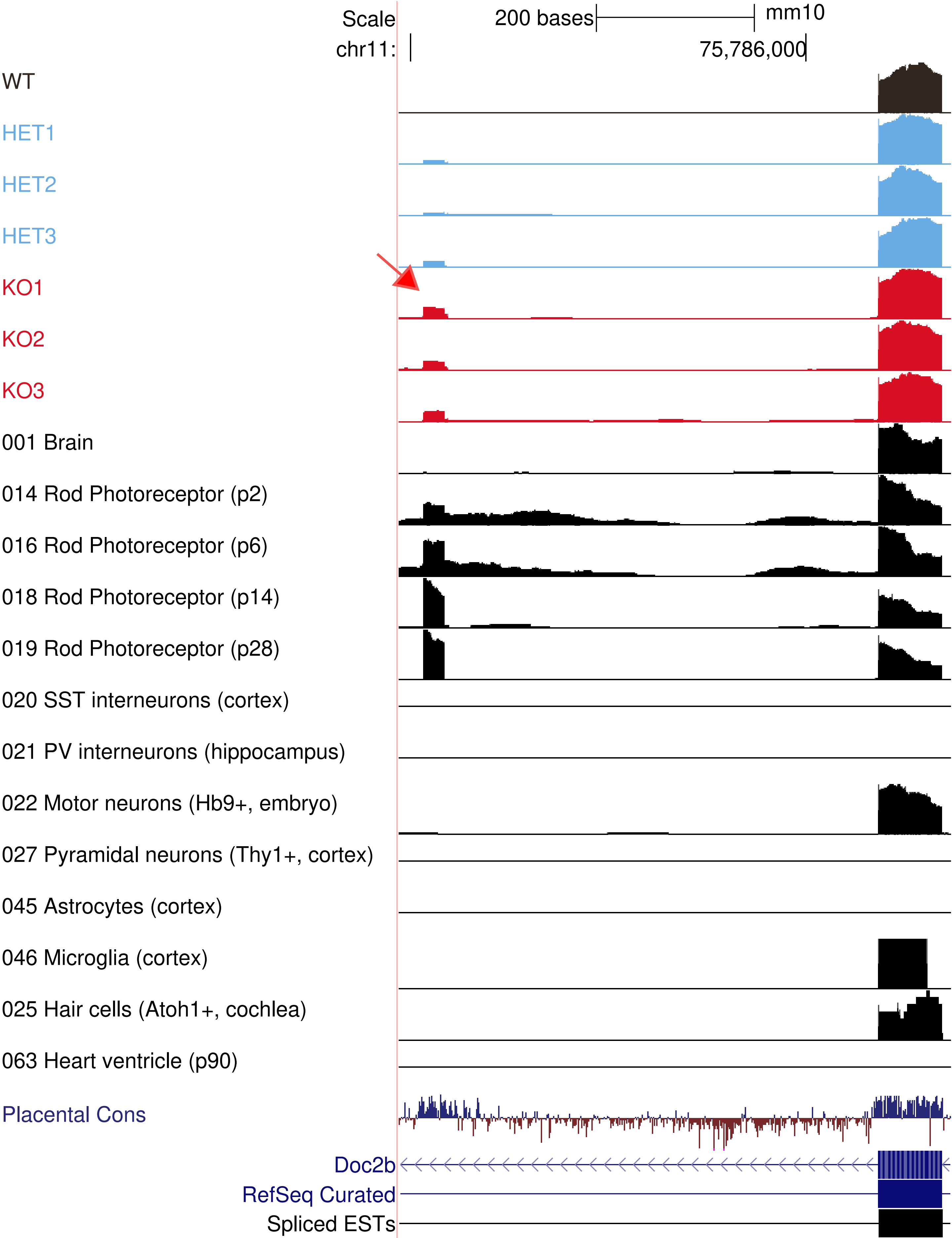

Scale
200 bases
mm10
chr11:
75,786,000
WT
HET1
HET2
HET3
KO1
KO2
KO3
001 Brain
014 Rod Photoreceptor (p2)
016 Rod Photoreceptor (p6)
018 Rod Photoreceptor (p14)
019 Rod Photoreceptor (p28)
020 SST interneurons (cortex)
021 PV interneurons (hippocampus)
022 Motor neurons (Hb9+, embryo)
027 Pyramidal neurons (Thy1+, cortex)
045 Astrocytes (cortex)
046 Microglia (cortex)
025 Hair cells (Atoh1+, cochlea)
063 Heart ventricle (p90)
Placental Cons
Doc2b
RefSeq Curated
Spliced ESTs

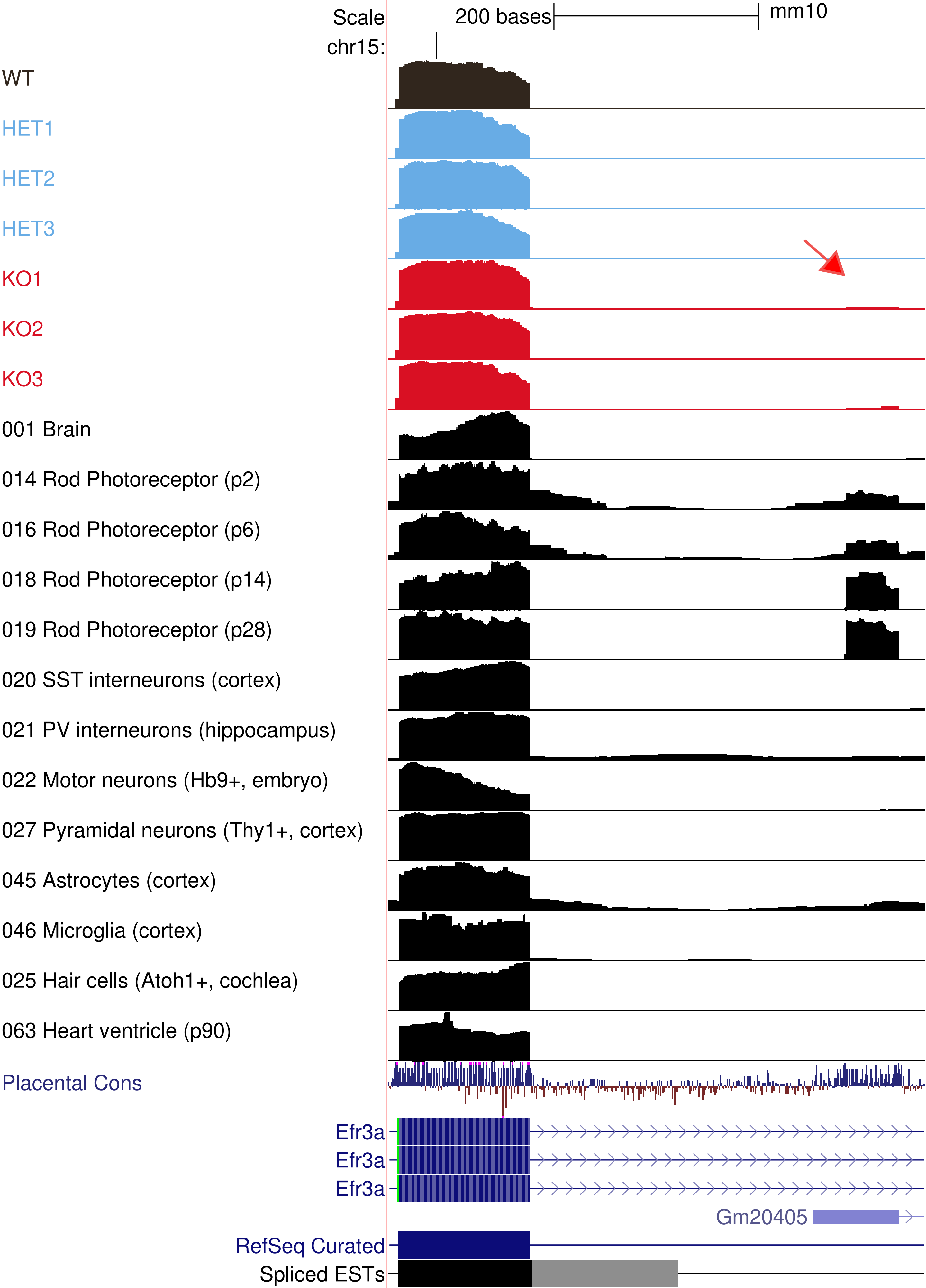
Scale
200 bases
mm10
chr15:
WT
HET1
HET2
HET3
KO1
KO2
KO3
001 Brain
014 Rod Photoreceptor (p2)
016 Rod Photoreceptor (p6)
018 Rod Photoreceptor (p14)
019 Rod Photoreceptor (p28)
020 SST interneurons (cortex)
021 PV interneurons (hippocampus)
022 Motor neurons (Hb9+, embryo)
027 Pyramidal neurons (Thy1+, cortex)
045 Astrocytes (cortex)
046 Microglia (cortex)
025 Hair cells (Atoh1+, cochlea)
063 Heart ventricle (p90)
Placental Cons
Efr3a
Efr3a
Efr3a
Gm20405
RefSeq Curated
Spliced ESTs

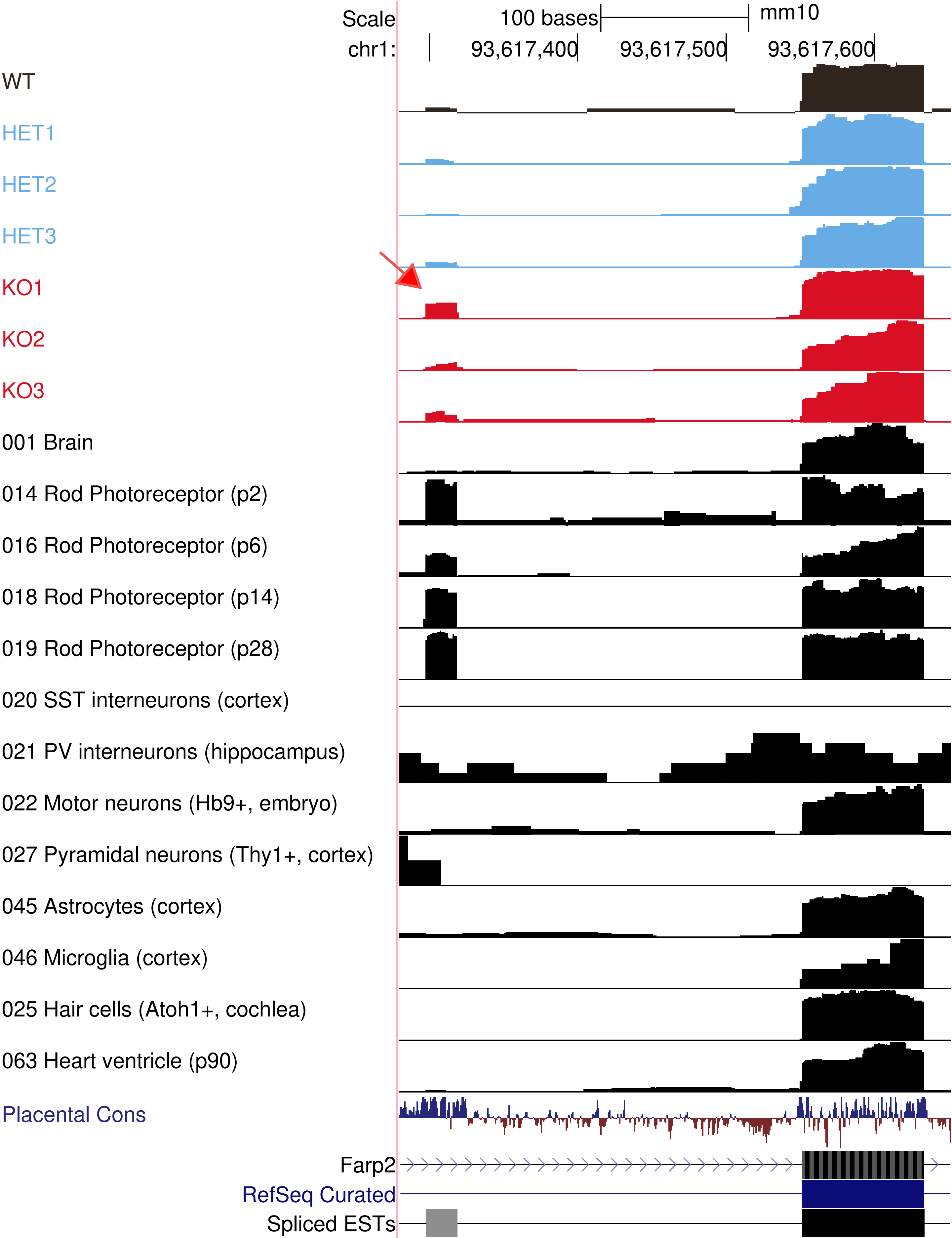

Scale
100 bases
mm10
chr1:
93,617,400
93,617,500
93,617,600
WT
HET1
HET2
HET3
KO1
KO2
KO3
001 Brain
014 Rod Photoreceptor (p2)
016 Rod Photoreceptor (p6)
018 Rod Photoreceptor (p14)
019 Rod Photoreceptor (p28)
020 SST interneurons (cortex)
021 PV interneurons (hippocampus)
022 Motor neurons (Hb9+, embryo)
027 Pyramidal neurons (Thy1+, cortex)
045 Astrocytes (cortex)
046 Microglia (cortex)
025 Hair cells (Atoh1+, cochlea)
063 Heart ventricle (p90)
Placental Cons
Farp2
RefSeq Curated
Spliced ESTs

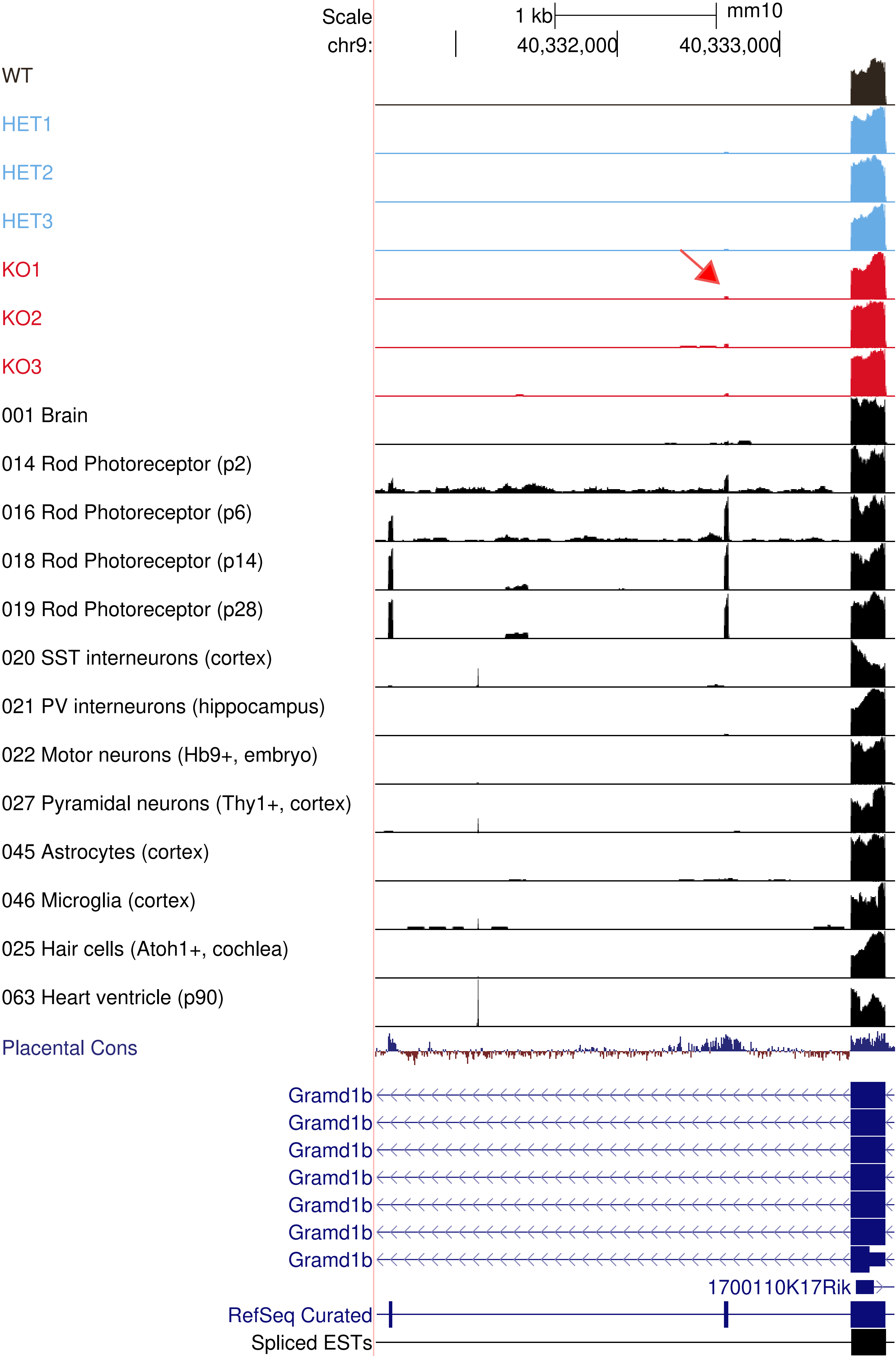

Scale
1 kb
mm10
chr9:
40,332,000
40,333,000
WT
HET1
HET2
HET3
KO1
KO2
KO3
001 Brain
014 Rod Photoreceptor (p2)
016 Rod Photoreceptor (p6)
018 Rod Photoreceptor (p14)
019 Rod Photoreceptor (p28)
020 SST interneurons (cortex)
021 PV interneurons (hippocampus)
022 Motor neurons (Hb9+, embryo)
027 Pyramidal neurons (Thy1+, cortex)
045 Astrocytes (cortex)
046 Microglia (cortex)
025 Hair cells (Atoh1+, cochlea)
063 Heart ventricle (p90)
Placental Cons
Gramd1b
Gramd1b
Gramd1b
Gramd1b
Gramd1b
Gramd1b
Gramd1b
1700110K17Rik
RefSeq Curated
Spliced ESTs

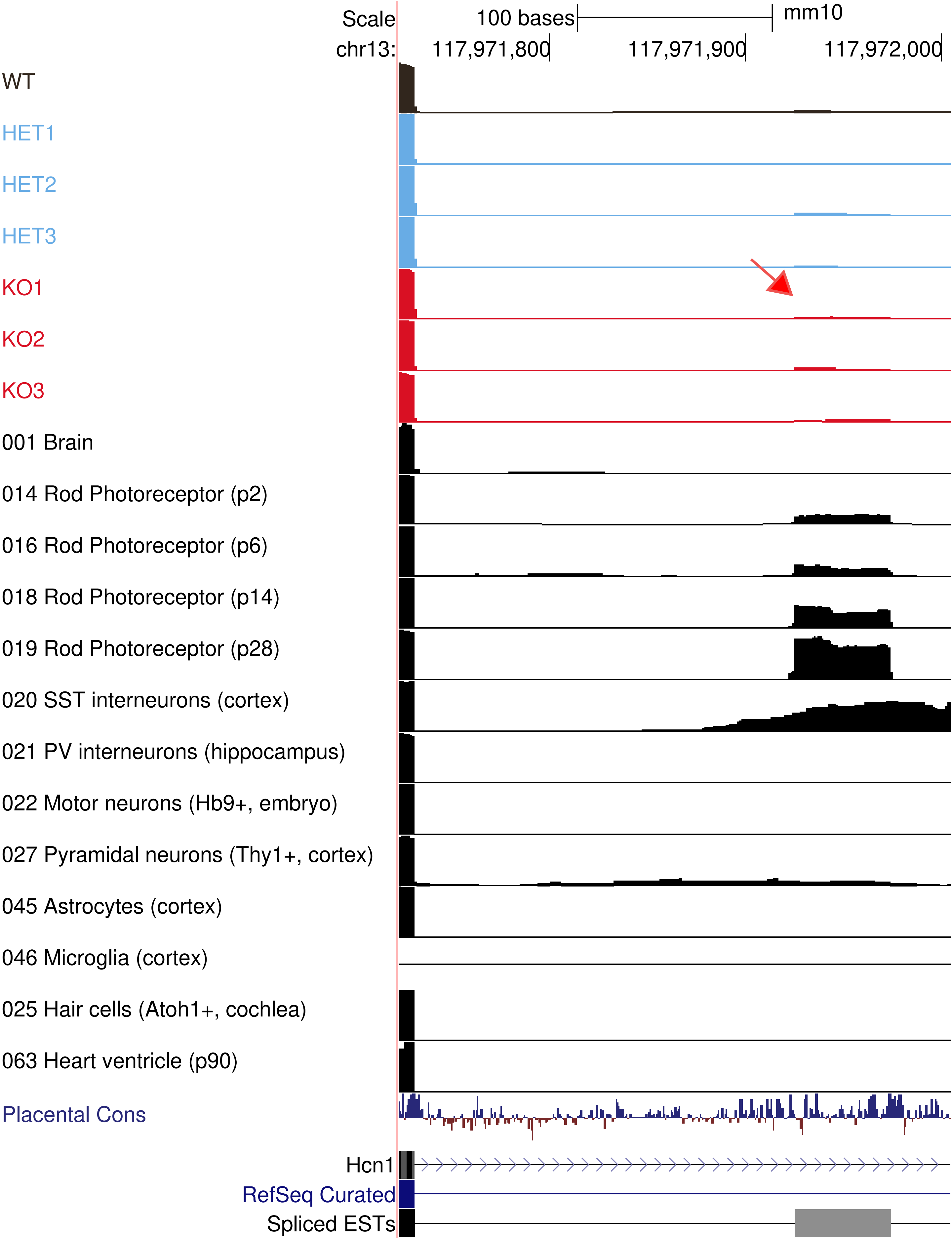
Scale
100 bases
mm10
chr13:
117,971,800
117,971,900
117,972,000
WT
HET1
HET2
HET3
KO1
KO2
KO3
001 Brain
014 Rod Photoreceptor (p2)
016 Rod Photoreceptor (p6)
018 Rod Photoreceptor (p14)
019 Rod Photoreceptor (p28)
020 SST interneurons (cortex)
021 PV interneurons (hippocampus)
022 Motor neurons (Hb9+, embryo)
027 Pyramidal neurons (Thy1+, cortex)
045 Astrocytes (cortex)
046 Microglia (cortex)
025 Hair cells (Atoh1+, cochlea)
063 Heart ventricle (p90)
Placental Cons
Hcn1
RefSeq Curated
Spliced ESTs

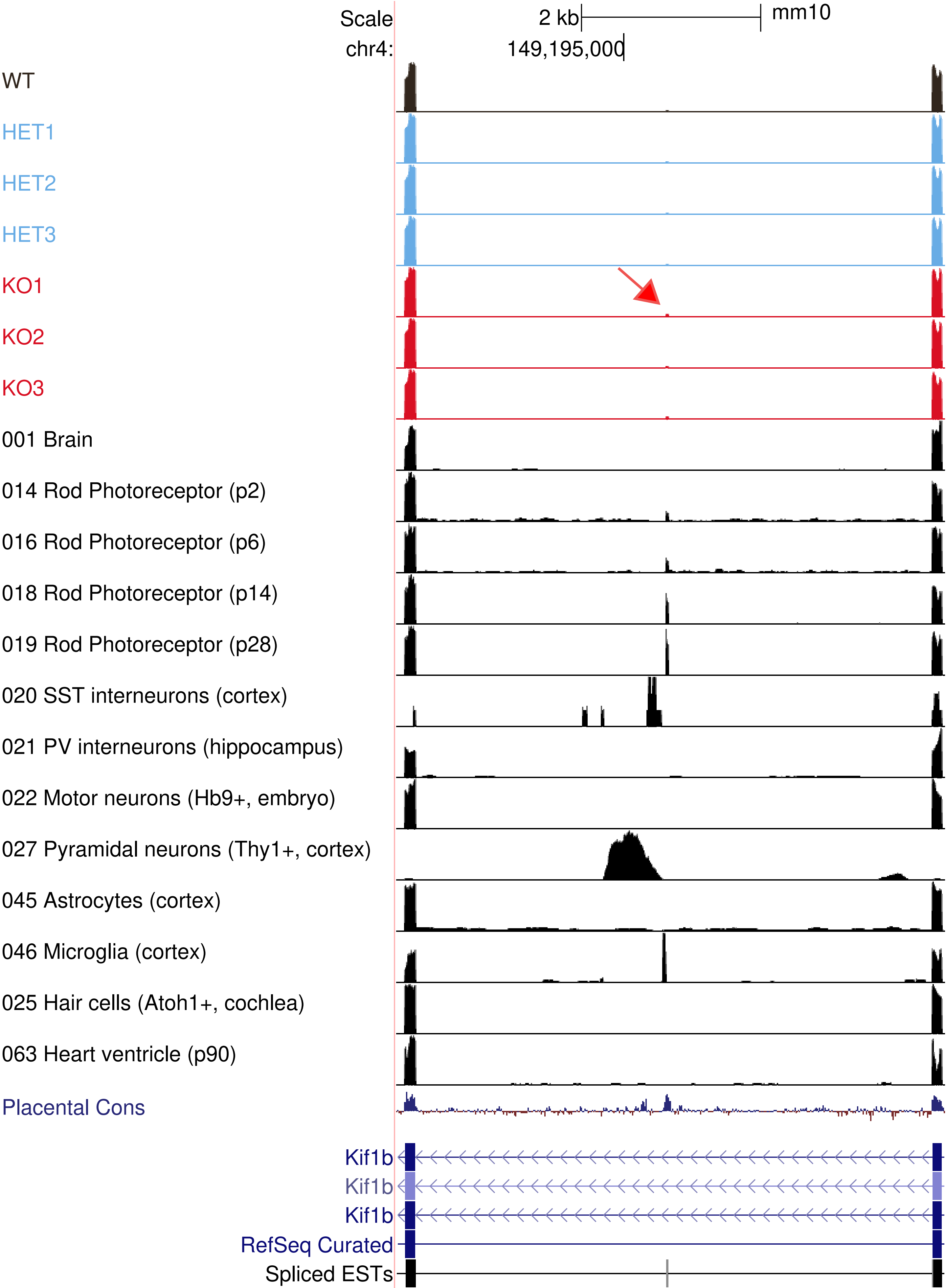
Scale
2 kb
mm10
chr4:
149,195,000
WT
HET1
HET2
HET3
KO1
KO2
KO3
001 Brain
014 Rod Photoreceptor (p2)
016 Rod Photoreceptor (p6)
018 Rod Photoreceptor (p14)
019 Rod Photoreceptor (p28)
020 SST interneurons (cortex)
021 PV interneurons (hippocampus)
022 Motor neurons (Hb9+, embryo)
027 Pyramidal neurons (Thy1+, cortex)
045 Astrocytes (cortex)
046 Microglia (cortex)
025 Hair cells (Atoh1+, cochlea)
063 Heart ventricle (p90)
Placental Cons
Kif1b
Kif1b
Kif1b
RefSeq Curated
Spliced ESTs

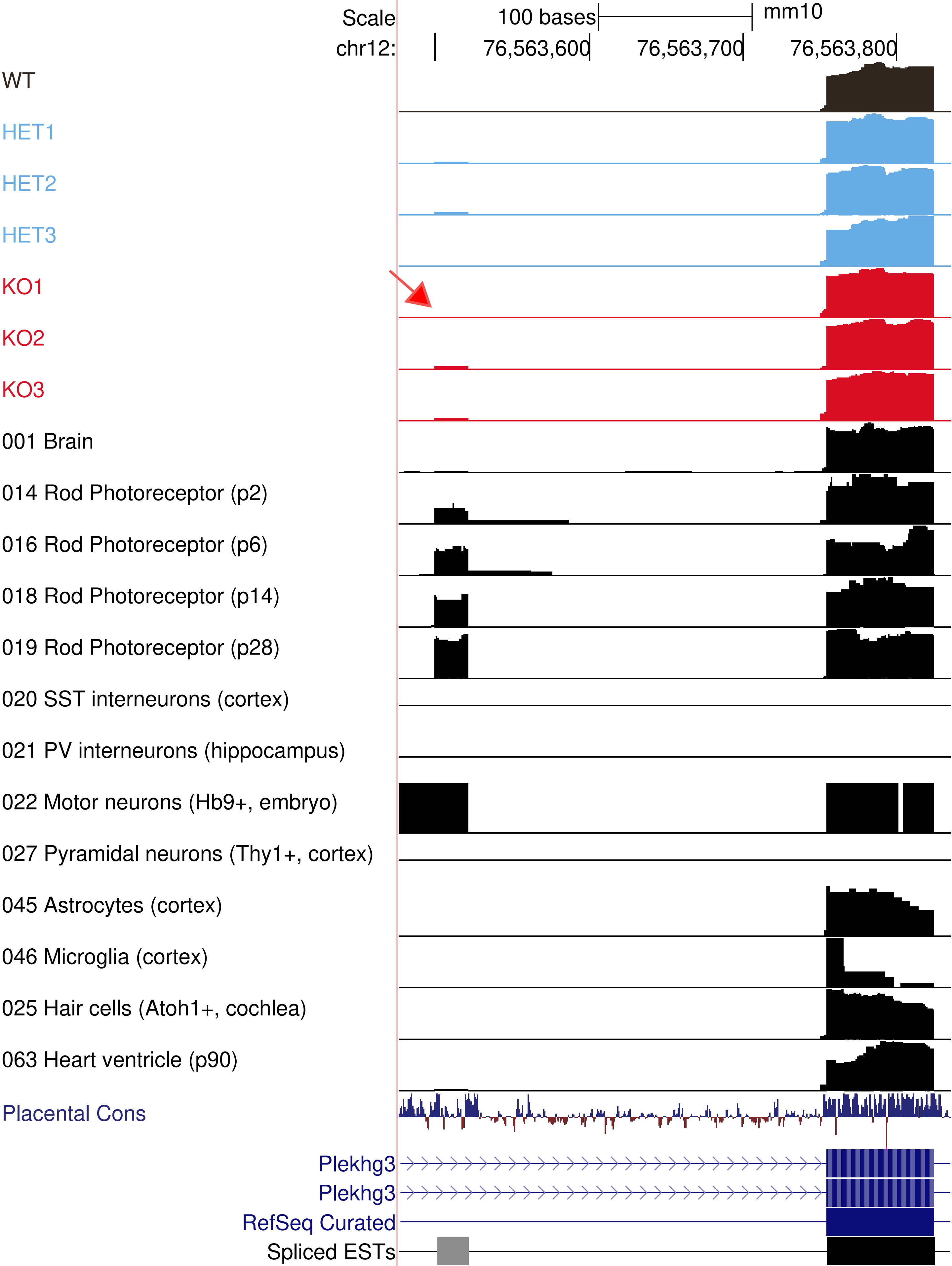

Scale
100 bases
mm10
chr12:
76,563,600
76,563,700
76,563,800
WT
HET1
HET2
HET3
KO1
KO2
KO3
001 Brain
014 Rod Photoreceptor (p2)
016 Rod Photoreceptor (p6)
018 Rod Photoreceptor (p14)
019 Rod Photoreceptor (p28)
020 SST interneurons (cortex)
021 PV interneurons (hippocampus)
022 Motor neurons (Hb9+, embryo)
027 Pyramidal neurons (Thy1+, cortex)
045 Astrocytes (cortex)
046 Microglia (cortex)
025 Hair cells (Atoh1+, cochlea)
063 Heart ventricle (p90)
Placental Cons
Plekhg3
Plekhg3
RefSeq Curated
Spliced ESTs

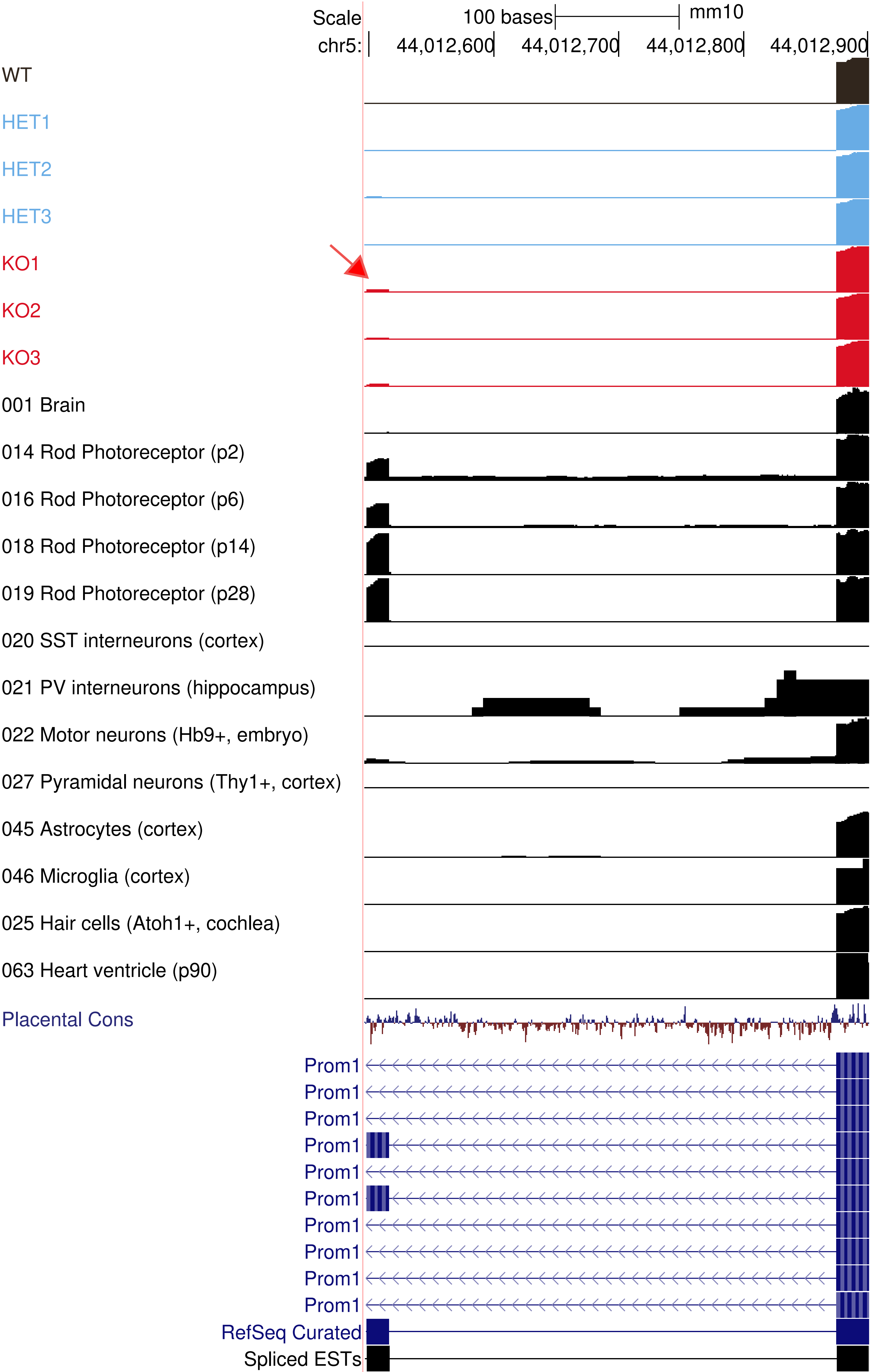
Scale
100 bases
mm10
chr5:
44,012,600
44,012,700
44,012,800
44,012,900
WT
HET1
HET2
HET3
KO1
KO2
KO3
001 Brain
014 Rod Photoreceptor (p2)
016 Rod Photoreceptor (p6)
018 Rod Photoreceptor (p14)
019 Rod Photoreceptor (p28)
020 SST interneurons (cortex)
021 PV interneurons (hippocampus)
022 Motor neurons (Hb9+, embryo)
027 Pyramidal neurons (Thy1+, cortex)
045 Astrocytes (cortex)
046 Microglia (cortex)
025 Hair cells (Atoh1+, cochlea)
063 Heart ventricle (p90)
Placental Cons
Prom1
Prom1
Prom1
Prom1
Prom1
Prom1
Prom1
Prom1
Prom1
Prom1
RefSeq Curated
Spliced ESTs

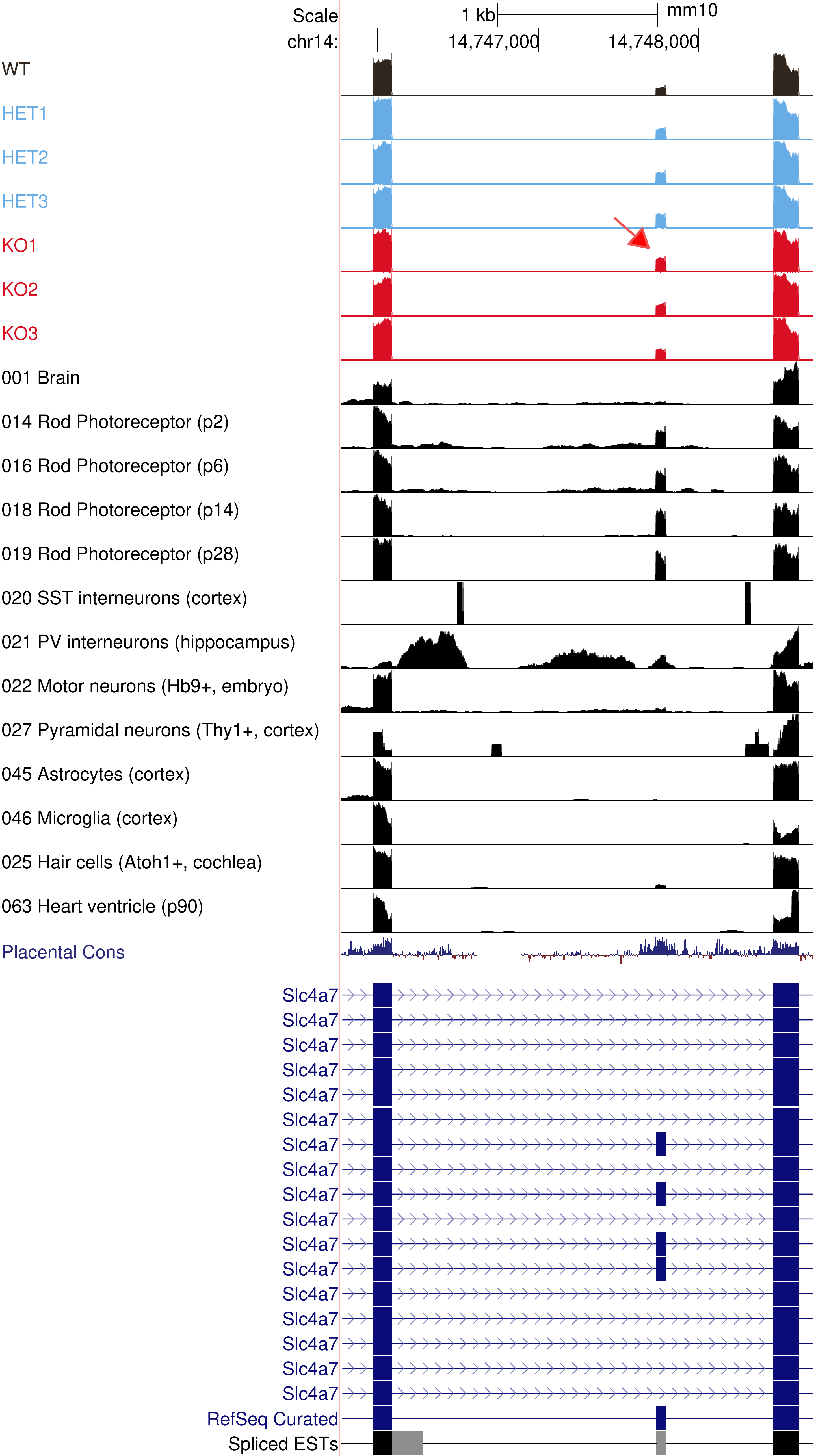
Scale
1 kb
mm10
chr14:
14,747,000
14,748,000
WT
HET1
HET2
HET3
KO1
KO2
KO3
001 Brain
014 Rod Photoreceptor (p2)
016 Rod Photoreceptor (p6)
018 Rod Photoreceptor (p14)
019 Rod Photoreceptor (p28)
020 SST interneurons (cortex)
021 PV interneurons (hippocampus)
022 Motor neurons (Hb9+, embryo)
027 Pyramidal neurons (Thy1+, cortex)
045 Astrocytes (cortex)
046 Microglia (cortex)
025 Hair cells (Atoh1+, cochlea)
063 Heart ventricle (p90)
Placental Cons
Slc4a7
Slc4a7
Slc4a7
Slc4a7
Slc4a7
Slc4a7
Slc4a7
Slc4a7
Slc4a7
Slc4a7
Slc4a7
Slc4a7
Slc4a7
Slc4a7
Slc4a7
Slc4a7
Slc4a7
RefSeq Curated
Spliced ESTs

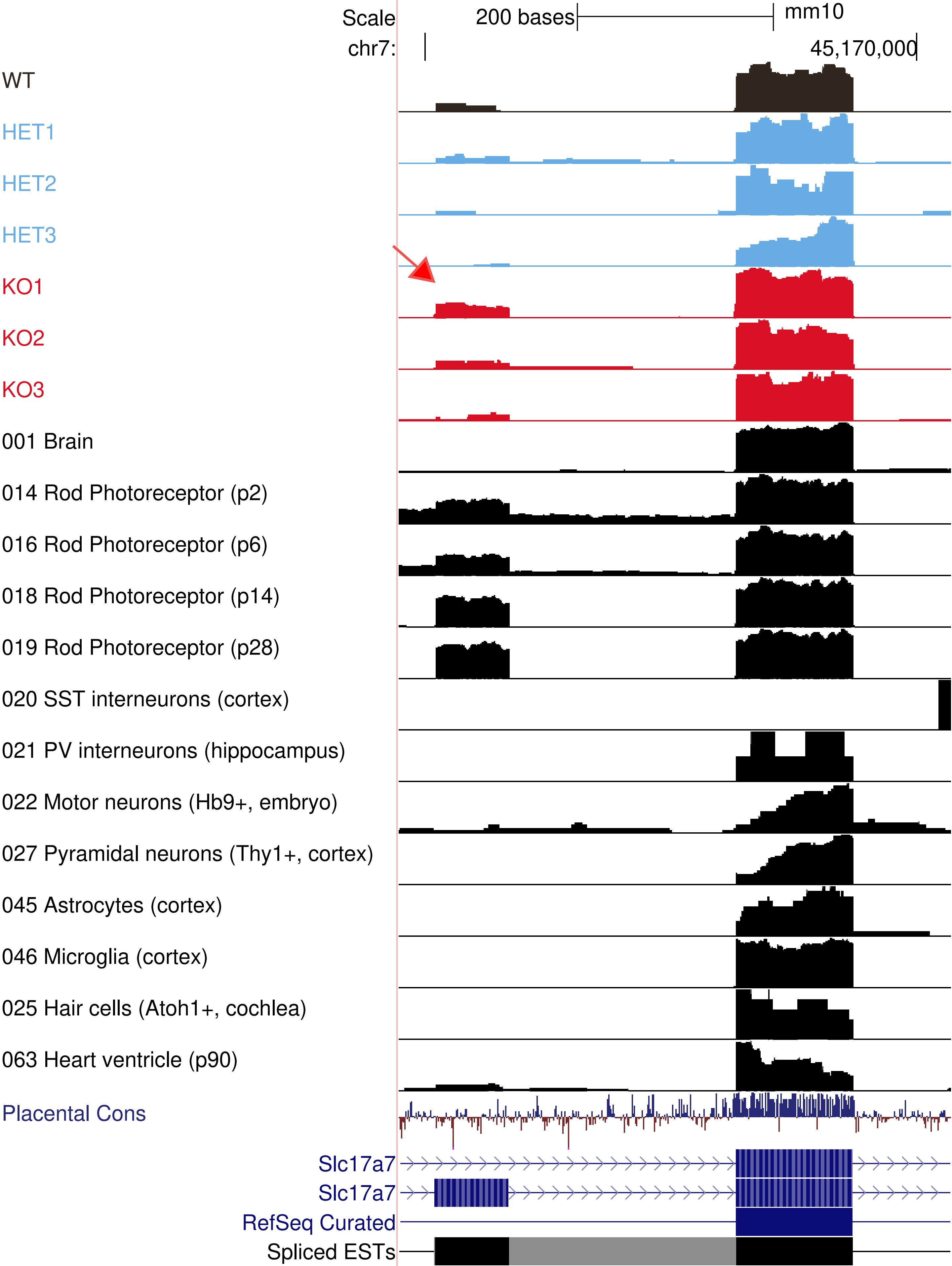

Scale
200 bases
mm10
chr7:
45,170,000
WT
HET1
HET2
HET3
KO1
KO2
KO3
001 Brain
014 Rod Photoreceptor (p2)
016 Rod Photoreceptor (p6)
018 Rod Photoreceptor (p14)
019 Rod Photoreceptor (p28)
020 SST interneurons (cortex)
021 PV interneurons (hippocampus)
022 Motor neurons (Hb9+, embryo)
027 Pyramidal neurons (Thy1+, cortex)
045 Astrocytes (cortex)
046 Microglia (cortex)
025 Hair cells (Atoh1+, cochlea)
063 Heart ventricle (p90)
Placental Cons
Slc17a7
Slc17a7
RefSeq Curated
Spliced ESTs

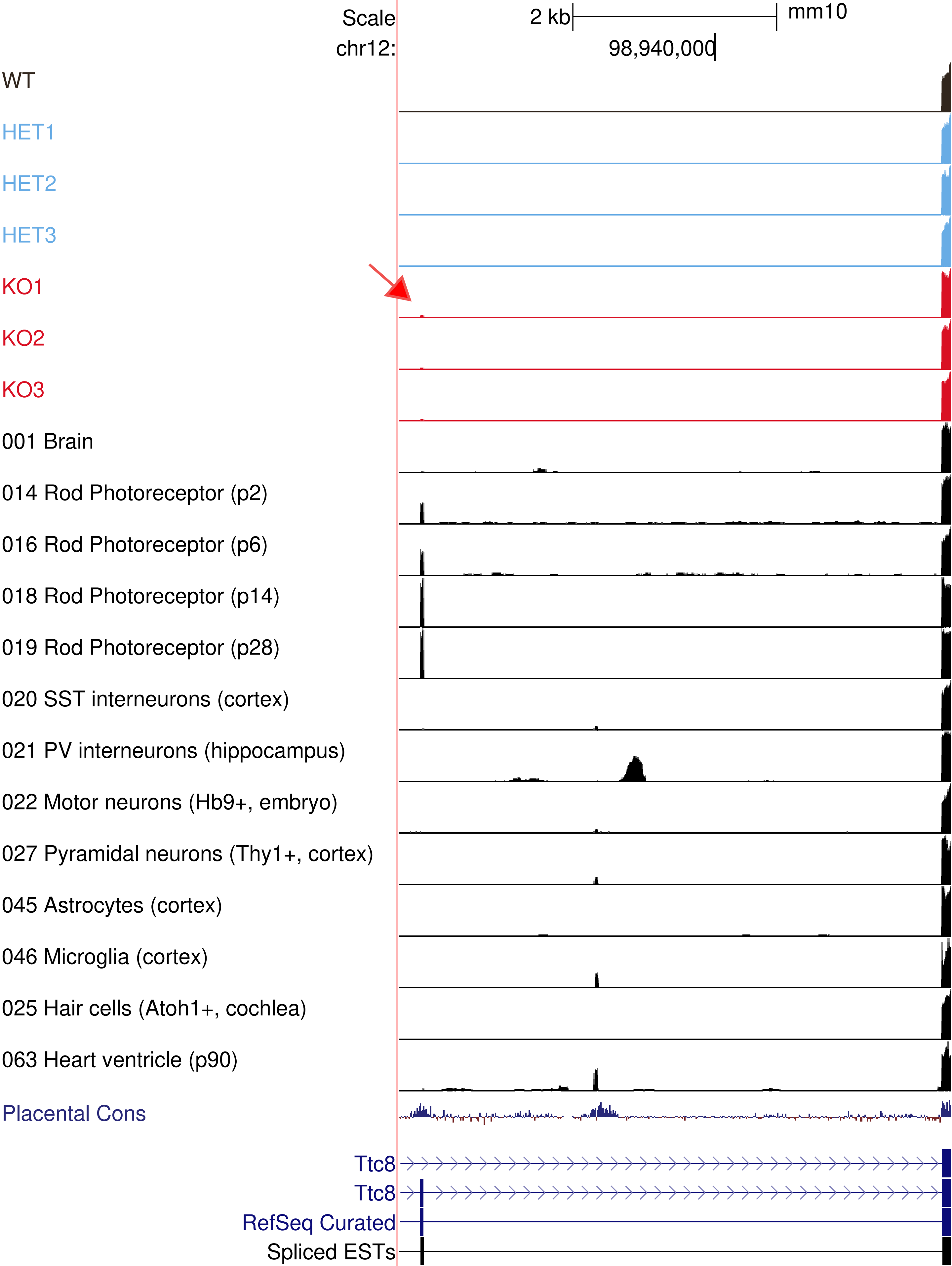
Scale
2 kb
mm10
chr12:
98,940,000
WT
HET1
HET2
HET3
KO1
KO2
KO3
001 Brain
014 Rod Photoreceptor (p2)
016 Rod Photoreceptor (p6)
018 Rod Photoreceptor (p14)
019 Rod Photoreceptor (p28)
020 SST interneurons (cortex)
021 PV interneurons (hippocampus)
022 Motor neurons (Hb9+, embryo)
027 Pyramidal neurons (Thy1+, cortex)
045 Astrocytes (cortex)
046 Microglia (cortex)
025 Hair cells (Atoh1+, cochlea)
063 Heart ventricle (p90)
Placental Cons
Ttc8
Ttc8
RefSeq Curated
Spliced ESTs

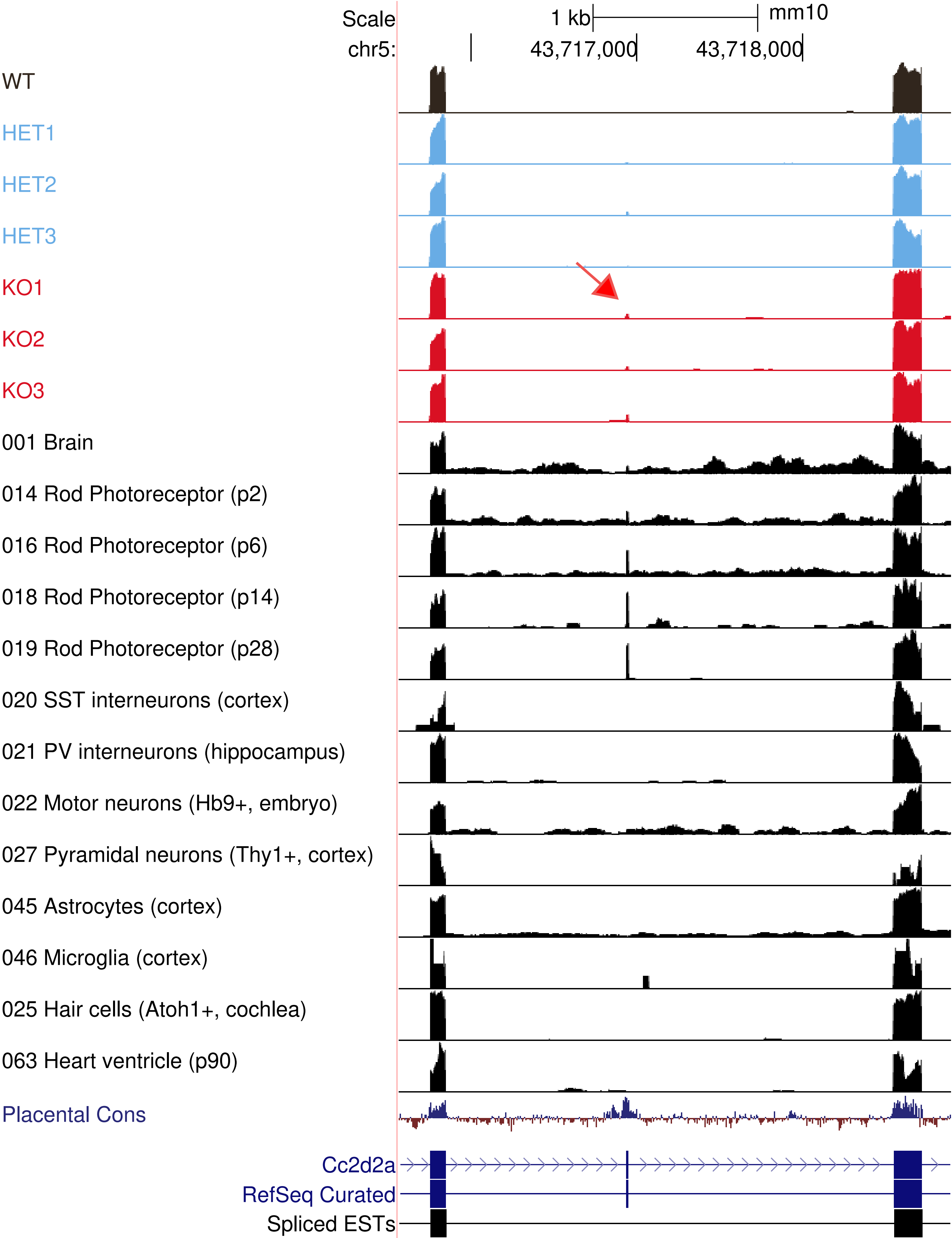
Scale
1 kb
mm10
chr5:
43,717,000
43,718,000
WT
HET1
HET2
HET3
KO1
KO2
KO3
001 Brain
014 Rod Photoreceptor (p2)
016 Rod Photoreceptor (p6)
018 Rod Photoreceptor (p14)
019 Rod Photoreceptor (p28)
020 SST interneurons (cortex)
021 PV interneurons (hippocampus)
022 Motor neurons (Hb9+, embryo)
027 Pyramidal neurons (Thy1+, cortex)
045 Astrocytes (cortex)
046 Microglia (cortex)
025 Hair cells (Atoh1+, cochlea)
063 Heart ventricle (p90)
Placental Cons
Cc2d2a
RefSeq Curated
Spliced ESTs

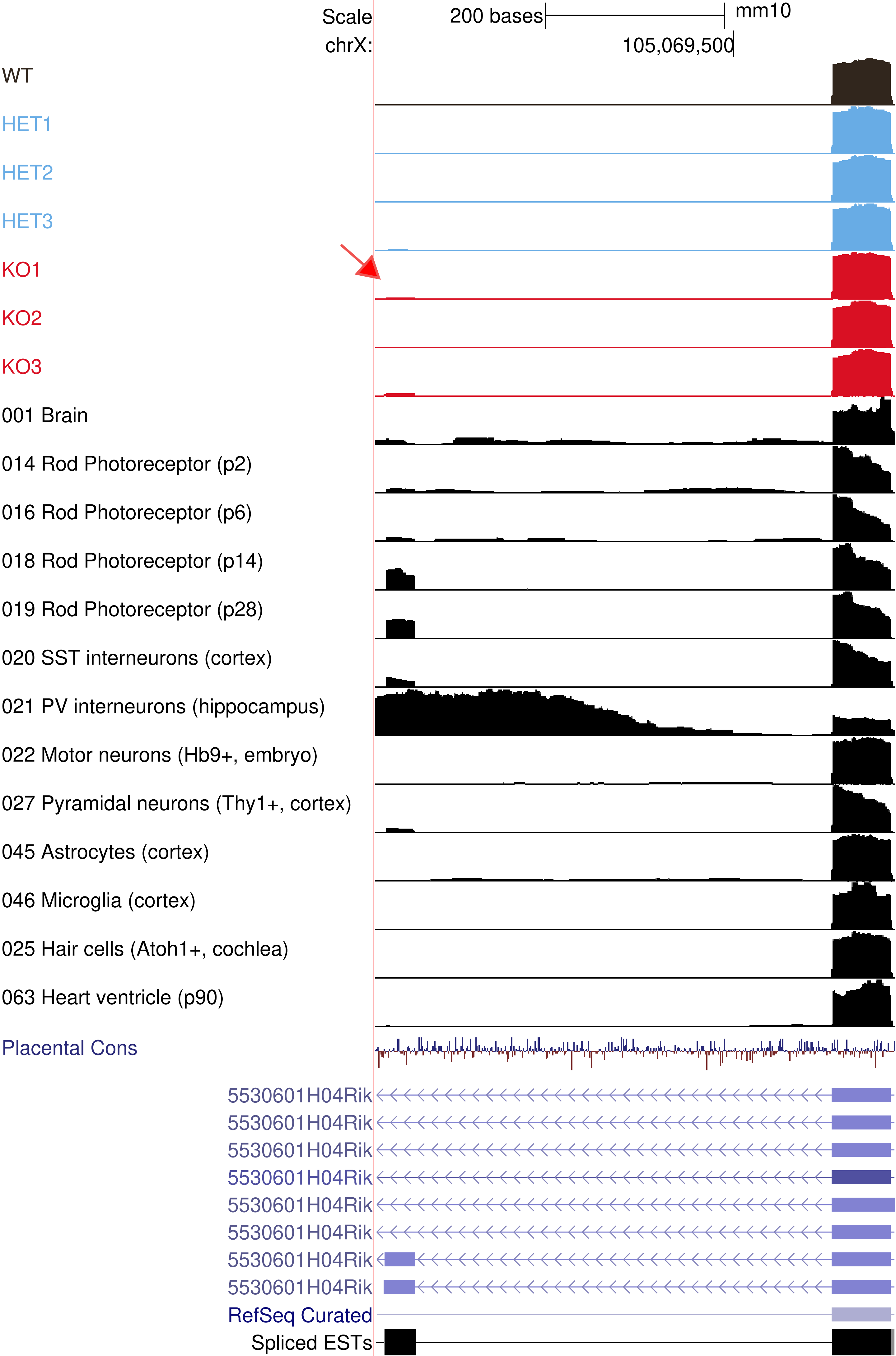

Scale
200 bases
mm10
chrX:
105,069,500
WT
HET1
HET2
HET3
KO1
KO2
KO3
001 Brain
014 Rod Photoreceptor (p2)
016 Rod Photoreceptor (p6)
018 Rod Photoreceptor (p14)
019 Rod Photoreceptor (p28)
020 SST interneurons (cortex)
021 PV interneurons (hippocampus)
022 Motor neurons (Hb9+, embryo)
027 Pyramidal neurons (Thy1+, cortex)
045 Astrocytes (cortex)
046 Microglia (cortex)
025 Hair cells (Atoh1+, cochlea)
063 Heart ventricle (p90)
Placental Cons
5530601H04Rik
5530601H04Rik
5530601H04Rik
5530601H04Rik
5530601H04Rik
5530601H04Rik
5530601H04Rik
5530601H04Rik
RefSeq Curated
Spliced ESTs

Extended Dataset 3

B. Neuron Specific Exons

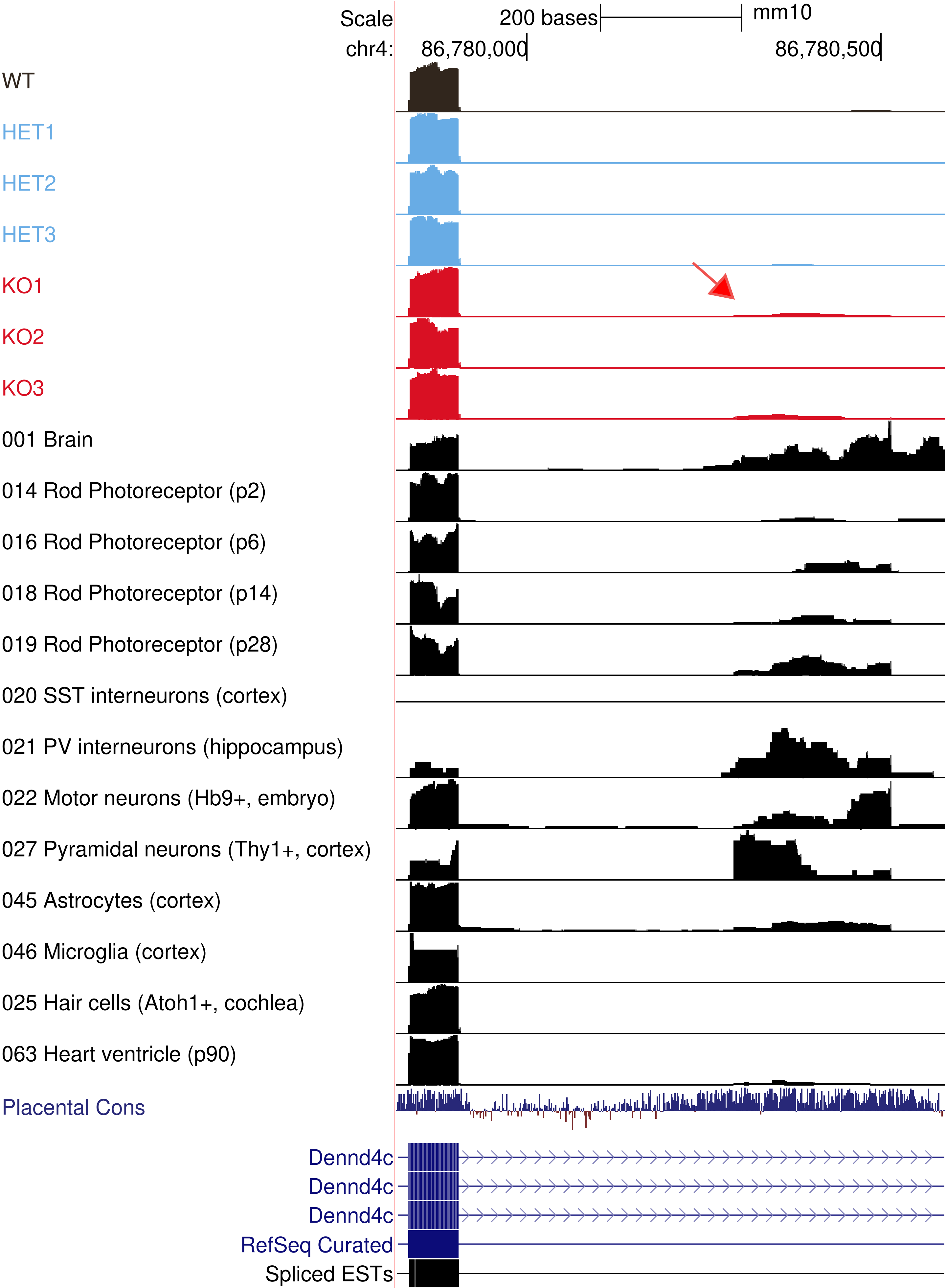

Scale
200 bases
mm10
chr4:
86,780,000
86,780,500
WT
HET1
HET2
HET3
KO1
KO2
KO3
001 Brain
014 Rod Photoreceptor (p2)
016 Rod Photoreceptor (p6)
018 Rod Photoreceptor (p14)
019 Rod Photoreceptor (p28)
020 SST interneurons (cortex)
021 PV interneurons (hippocampus)
022 Motor neurons (Hb9+, embryo)
027 Pyramidal neurons (Thy1+, cortex)
045 Astrocytes (cortex)
046 Microglia (cortex)
025 Hair cells (Atoh1+, cochlea)
063 Heart ventricle (p90)
Placental Cons
Dennd4c
Dennd4c
Dennd4c
RefSeq Curated
Spliced ESTs

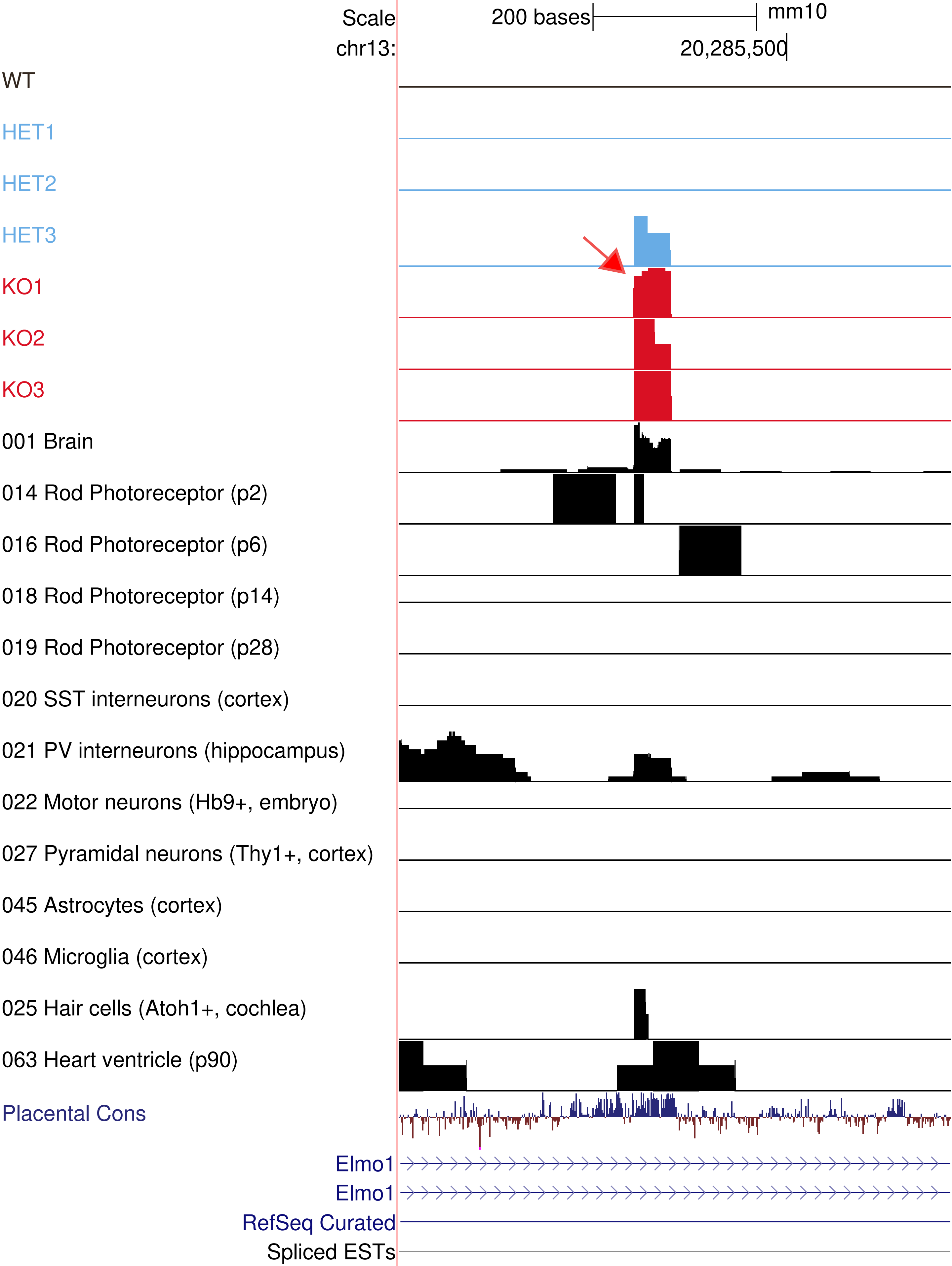
Scale
chr13:
200 bases
mm10
20,285,500
WT
HET1
HET2
HET3
KO1
KO2
KO3
001 Brain
014 Rod Photoreceptor (p2)
016 Rod Photoreceptor (p6)
018 Rod Photoreceptor (p14)
019 Rod Photoreceptor (p28)
020 SST interneurons (cortex)
021 PV interneurons (hippocampus)
022 Motor neurons (Hb9+, embryo)
027 Pyramidal neurons (Thy1+, cortex)
045 Astrocytes (cortex)
046 Microglia (cortex)
025 Hair cells (Atoh1+, cochlea)
063 Heart ventricle (p90)
Placental Cons
Elmo1
Elmo1
RefSeq Curated
Spliced ESTs

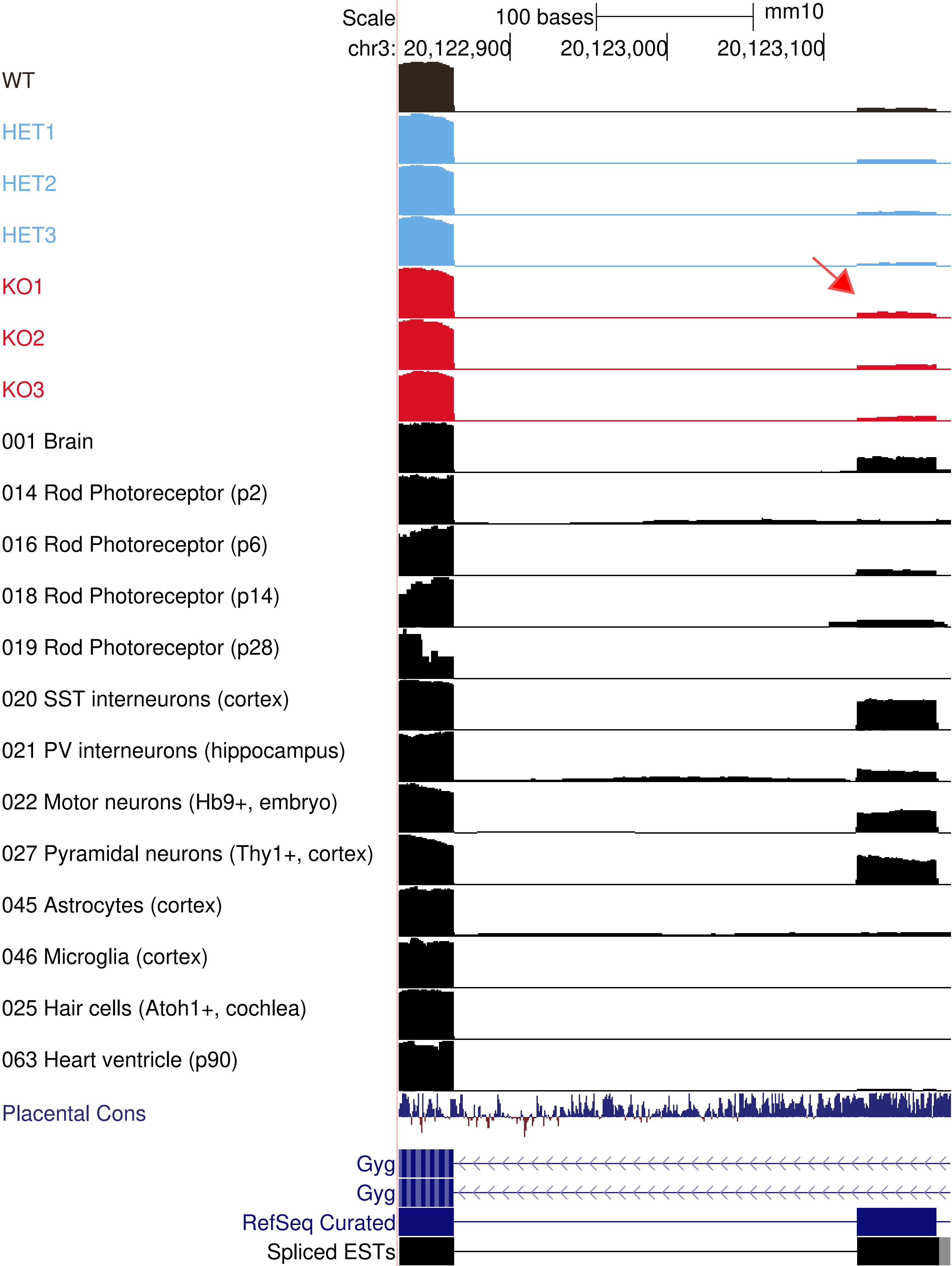
Scale
100 bases
mm10
chr3: 20,122,900
20,123,000
20,123,100
WT
HET1
HET2
HET3
KO1
KO2
KO3
001 Brain
014 Rod Photoreceptor (p2)
016 Rod Photoreceptor (p6)
018 Rod Photoreceptor (p14)
019 Rod Photoreceptor (p28)
020 SST interneurons (cortex)
021 PV interneurons (hippocampus)
022 Motor neurons (Hb9+, embryo)
027 Pyramidal neurons (Thy1+, cortex)
045 Astrocytes (cortex)
046 Microglia (cortex)
025 Hair cells (Atoh1+, cochlea)
063 Heart ventricle (p90)
Placental Cons
Gyg
Gyg
RefSeq Curated
Spliced ESTs

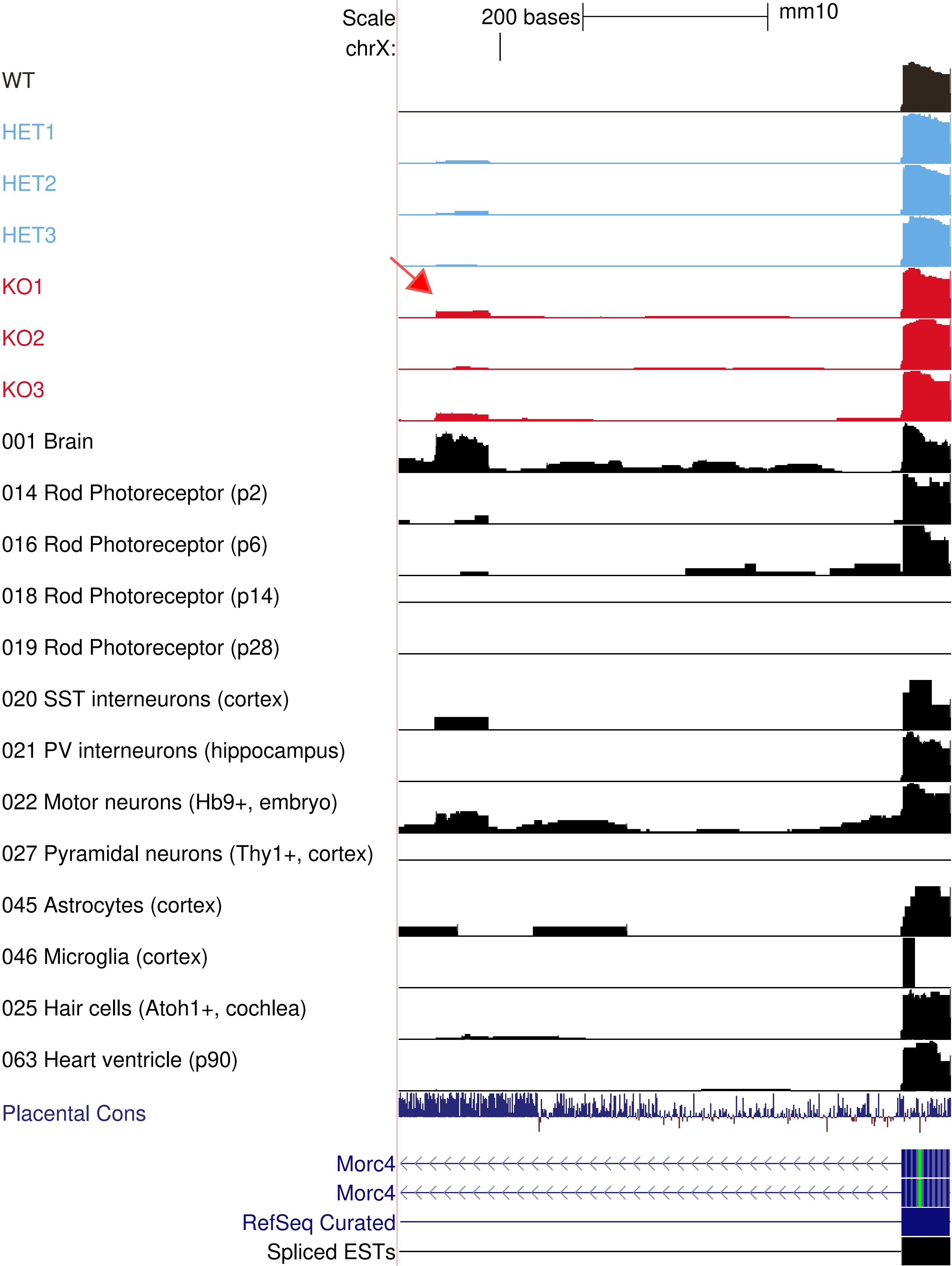

Scale
200 bases
mm10
chrX:
WT
HET1
HET2
HET3
KO1
KO2
KO3
001 Brain
014 Rod Photoreceptor (p2)
016 Rod Photoreceptor (p6)
018 Rod Photoreceptor (p14)
019 Rod Photoreceptor (p28)
020 SST interneurons (cortex)
021 PV interneurons (hippocampus)
022 Motor neurons (Hb9+, embryo)
027 Pyramidal neurons (Thy1+, cortex)
045 Astrocytes (cortex)
046 Microglia (cortex)
025 Hair cells (Atoh1+, cochlea)
063 Heart ventricle (p90)
Placental Cons
Morc4
Morc4
RefSeq Curated
Spliced ESTs

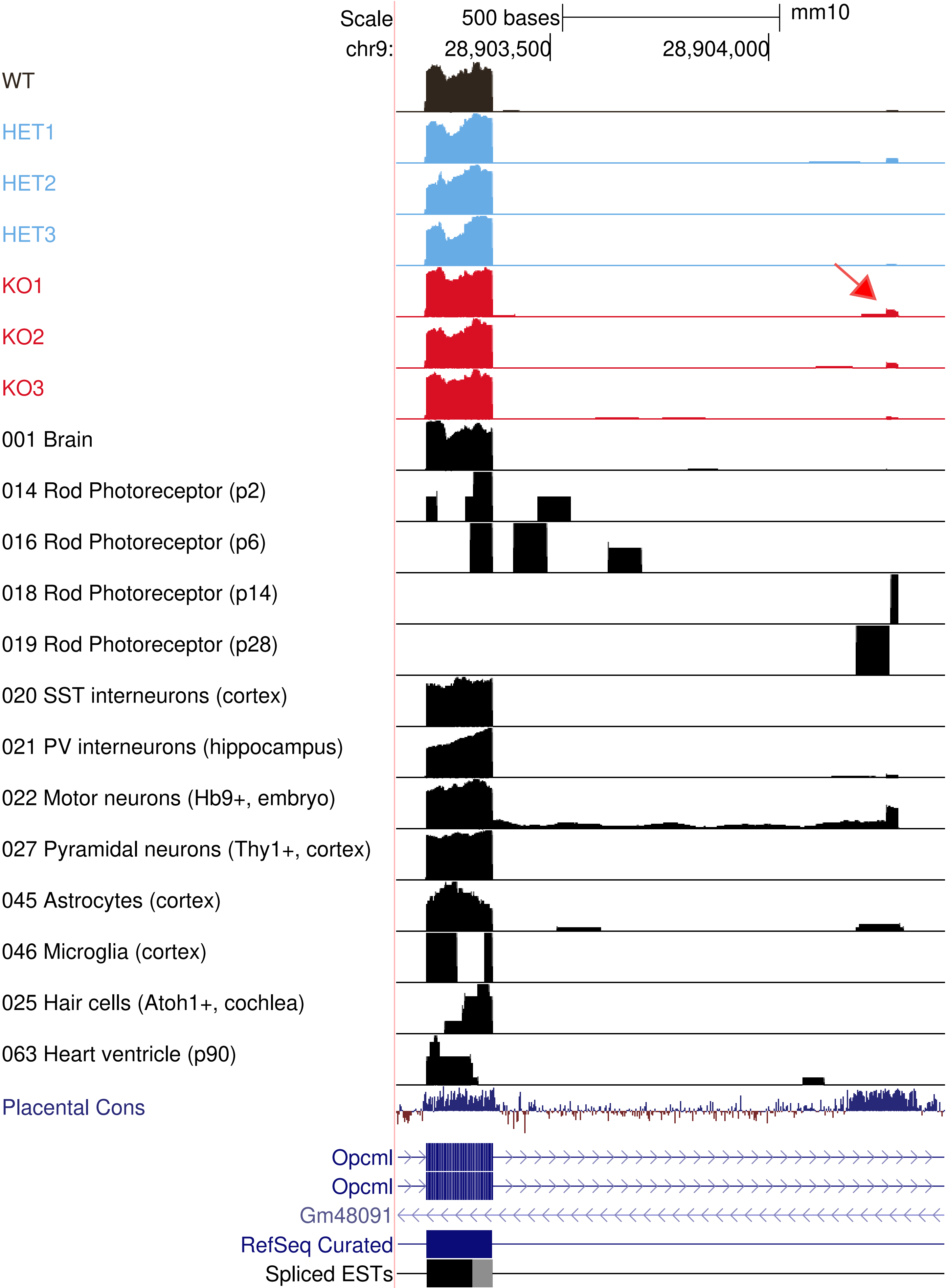

Scale
500 bases
mm10
chr9:
28,903,500
28,904,000
WT
HET1
HET2
HET3
KO1
KO2
KO3
001 Brain
014 Rod Photoreceptor (p2)
016 Rod Photoreceptor (p6)
018 Rod Photoreceptor (p14)
019 Rod Photoreceptor (p28)
020 SST interneurons (cortex)
021 PV interneurons (hippocampus)
022 Motor neurons (Hb9+, embryo)
027 Pyramidal neurons (Thy1+, cortex)
045 Astrocytes (cortex)
046 Microglia (cortex)
025 Hair cells (Atoh1+, cochlea)
063 Heart ventricle (p90)
Placental Cons
Opcml
Opcml
Gm48091
RefSeq Curated
Spliced ESTs

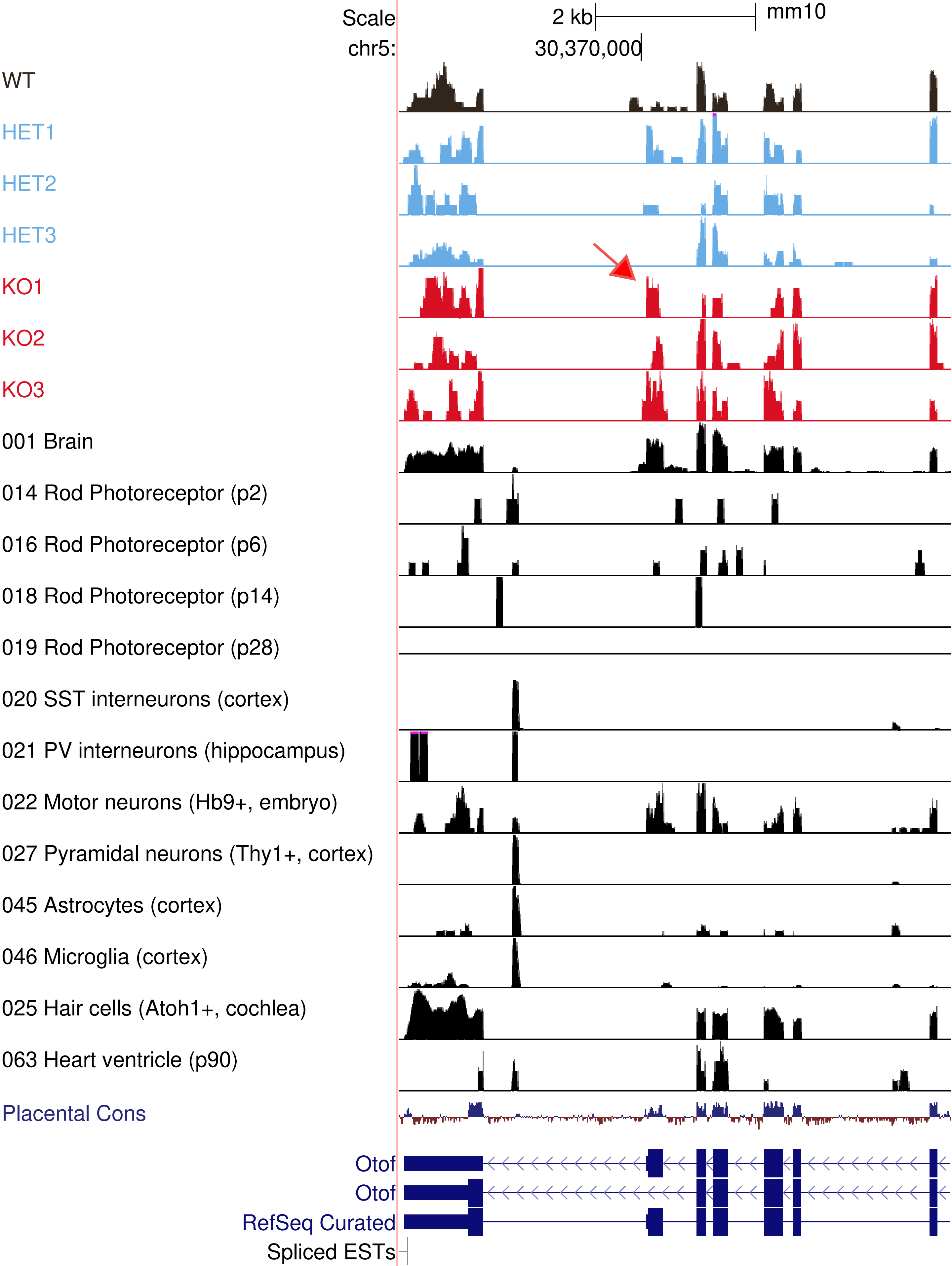
Scale
2 kb
mm10
chr5:
30,370,000
WT
HET1
HET2
HET3
KO1
KO2
KO3
001 Brain
014 Rod Photoreceptor (p2)
016 Rod Photoreceptor (p6)
018 Rod Photoreceptor (p14)
019 Rod Photoreceptor (p28)
020 SST interneurons (cortex)
021 PV interneurons (hippocampus)
022 Motor neurons (Hb9+, embryo)
027 Pyramidal neurons (Thy1+, cortex)
045 Astrocytes (cortex)
046 Microglia (cortex)
025 Hair cells (Atoh1+, cochlea)
063 Heart ventricle (p90)
Placental Cons
Otof
Otof
RefSeq Curated
Spliced ESTs

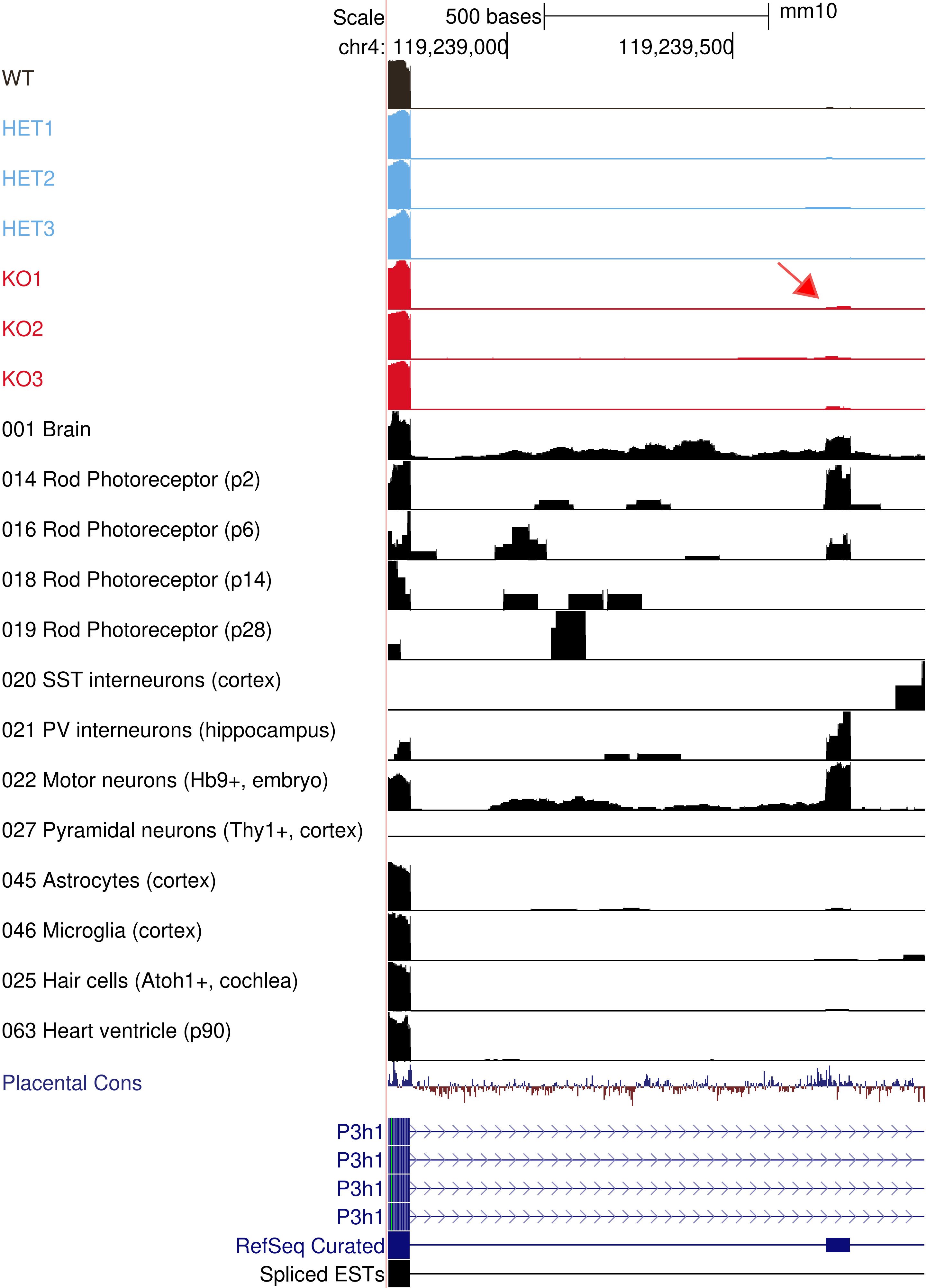

Scale
500 bases
mm10
chr4: 119,239,000
119,239,500
WT
HET1
HET2
HET3
KO1
KO2
KO3
001 Brain
014 Rod Photoreceptor (p2)
016 Rod Photoreceptor (p6)
018 Rod Photoreceptor (p14)
019 Rod Photoreceptor (p28)
020 SST interneurons (cortex)
021 PV interneurons (hippocampus)
022 Motor neurons (Hb9+, embryo)
027 Pyramidal neurons (Thy1+, cortex)
045 Astrocytes (cortex)
046 Microglia (cortex)
025 Hair cells (Atoh1+, cochlea)
063 Heart ventricle (p90)
Placental Cons
P3h1
P3h1
P3h1
P3h1
RefSeq Curated
Spliced ESTs

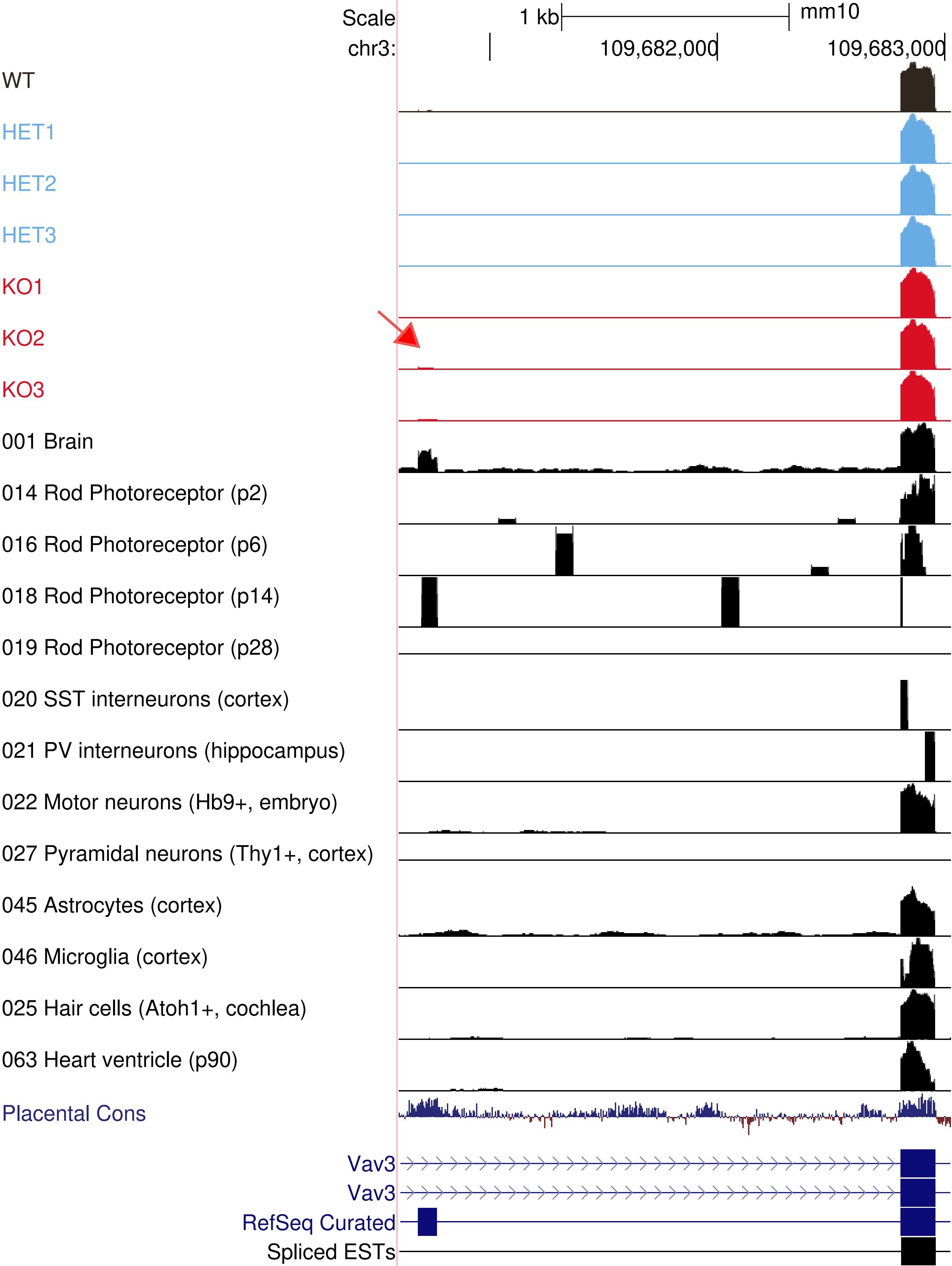
Scale
1 kb
mm10
chr3:
109,682,000
109,683,000
WT
HET1
HET2
HET3
KO1
KO2
KO3
001 Brain
014 Rod Photoreceptor (p2)
016 Rod Photoreceptor (p6)
018 Rod Photoreceptor (p14)
019 Rod Photoreceptor (p28)
020 SST interneurons (cortex)
021 PV interneurons (hippocampus)
022 Motor neurons (Hb9+, embryo)
027 Pyramidal neurons (Thy1+, cortex)
045 Astrocytes (cortex)
046 Microglia (cortex)
025 Hair cells (Atoh1+, cochlea)
063 Heart ventricle (p90)
Placental Cons
Vav3
Vav3
RefSeq Curated
Spliced ESTs
